# *Pparg* drives luminal differentiation and luminal tumor formation in the urothelium

**DOI:** 10.1101/2021.04.27.441646

**Authors:** Tiffany Tate, Tina Xiang, Mi Zhou, William Y. Kim, Xiao Chen, Hyunwoo Kim, Ekatherina Batourina, Chyuan-Sheng Lin, Chao Lu, Sara E. Wobker, James M. Mckiernan, Cathy Lee Mendelsohn

## Abstract

*Pparg*, a nuclear receptor, is downregulated in basal subtype bladder cancers that tend to be muscle invasive and amplified in luminal subtype bladder cancers that tend to be non-muscle invasive. Bladder cancers derive from the urothelium, one of the most quiescent epithelia in the body which is composed of basal, intermediate, and superficial cells. We find that expression of an activated form of Pparg (VP16;Pparg) in basal progenitors induces formation of superficial cells i*n situ*, that exit the cell cycle, and do not form tumors. Expression in basal progenitors that have been activated by mild injury however, results in luminal tumor formation. We find that tumors are immune deserted, which may be linked to downregulation of Nf-kb, a Pparg target. Interestingly, some luminal tumors begin to shift to basal subtype tumors with time, downregulating Pparg and other luminal markers. Our findings have important implications for treatment and diagnosis bladder cancer.

## Introduction

Bladder cancers are the 9th most common form of cancer worldwide, and the 4th most common among men (Sung et al, ACS 2021). Estimates from the CDC indicate that in 2021, there are about 83,730 new cases of bladder cancer (~64,280 in men and ~19,450 in women) in the United States and 17,200 deaths from bladder cancer (~12,260 men and ~4,940 women), with almost half (47%) of all cases attributable to smoking in the US (Society, 2020). Bladder cancer tumors arise from the urothelium, a water-proof barrier lining the urinary outflow tract that extends from the renal pelvis throughout the bladder and to the proximal urethra. The urothelium protects against infection, damage from toxins, and prevents exchange of fluids. It is one of the slowest cycling epithelia in the body (Jost, 1989), but undergoes a rapid sequence of exfoliation and regeneration in response to acute injury or infection. However, persistent or repeated damage and inflammation, can lead to permanent changes, including loss of endogenous urothelial populations and bladder pain disease (Birder, 2019; Ma et al., 2016) which has limited treatment options.

The urothelium is pseudostratified, containing two populations of basal cells; K14-Basal cells (K14+K5+) that are rare and reside exclusively in the basal layer and K5-Basal cells (K5+ K14-), which reside in the basal and suprabasal layers. Lineage studies suggest that K14-Basal cells are progenitors that can repopulate the urothelium *de novo* (Colopy et al., 2014; Kullmann et al., 2017; Narla et al., 2020; Schafer et al., 2017a; Wang et al., 2018), and are also cells of origin that can produce tumors (Papafotiou et al., 2016; Shin et al., 2011; Shin et al., 2014). Intermediate cells (I-cells) and Superficial cells (S-cells) which reside in upper urothelial layers, express luminal markers, including Krt20, Krt18, and Uroplakins (Upks). I-cells (P63+ UPKs+) are direct progenitors that replace S-cells when they die off (Colopy et al., 2014; Gandhi et al., 2013). They are attached to the basement membrane by a long cytoplasmic tail, and they can either divide or undergo failed cytokinesis producing binucleated I-cell (2n+2n) that undergo endoreplication, doubling DNA content without entering mitosis to produce 4n+4n S-cells (Wang et al., 2018). S-cells (UPK+ K20+ P63-) reside in the top layer; they are large, binucleated, and are critical for synthesis and transport of uroplakins, a family of proteins that assemble into tough crystalline plaques that line the apical surface of the urothelium. Although S-cells are post-mitotic, they are functionally very active since apical plaque is continuously degraded and replaced in response to stretch, as the bladder fills and empties (Apodaca, 2004; Carattino et al., 2013; Truschel et al., 2002; Wu et al., 1994; Yu et al., 2009).

Bladder cancers arise from the urothelium and were initially classified according to histological analysis prior to the availability of genomic sequencing including characteristics such as tumor grade and invasion into the muscularis propria (defining muscle invasive and non-muscle invasive lesions) (Guyon, 1884). 51% of diagnosed cases are non-muscle invasive bladder cancer (NMIBC) and 20-30% are muscle invasive bladder cancer [(MIBC)(Kamoun et al., 2020; SEER, 1969-2017)]. Approximately 10-20% of NMIBC cases eventually progress to MIBC (Kamoun et al., 2020; Knowles and Hurst, 2015; Tan et al., 2019). More recent studies from a number of groups have identified distinct subtypes of bladder cancer based on mutations and transcriptional profiles that cluster with the Luminal and Basal tumor types in MIBC. Approximately 47% of the cases are of the Luminal subtype and 35% are of the Basal subtype (Cancer Genome Atlas Research, 2014; Choi et al., 2014; Damrauer et al., 2014; Kamoun et al., 2020; Robertson et al., 2017; Sjodahl et al., 2012; Volkmer et al., 2012). These subtypes have grown in number and in general are encompassed in the Consensus Subtypes: luminal papillary (LumP), luminal nonspecified (LumNS), luminal unstable (LumU), stroma-rich, basal/squamous (Ba/Sq), and neuroendocrine-like [(NE-like)(Kamoun et al., 2020)]. Tumors with a basal/squamous subtype express a set of markers including KRT14, KRT5, CD44, and KRT6A. Basal/squamous tumors are generally immune infiltrated (Choi et al., 2014; Kamoun et al., 2020; Kardos et al., 2016; Rosenberg et al., 2016; Seiler et al., 2017). Bladder cancer of the luminal subtype express a set of markers found in urothelial I-cells and S-cells, including GATA3, FOXA1, and PPARG (Choi et al., 2014; Damrauer et al., 2014; Robertson et al., 2017; Sjödahl et al., 2014; Warrick et al., 2016). Luminal tumors are comparatively less invasive, but generally locally relapse after resection and also tend to be immune excluded (Kamoun et al., 2020; Kardos et al., 2016; Kim et al., 2019; Korpal et al., 2017; Saito et al., 2018).

PPARG-dependent transcription is important for a wide range of functions in different cell types, including adipogenesis, metabolism, and immunity [reviewed in:(Ahmadian et al., 2013; Lehrke and Lazar, 2005)]. PPARG agonists have profound effects on urothelial differentiation and *PPARG* mutations and amplifications contribute significantly to bladder cancers. *PPARG* is a member of the nuclear receptor superfamily of ligand-activated transcription factors that bind to response elements in regulatory regions of genes (Ahmadian et al., 2013; Barak et al., 1999). PPARG regulates transcription by forming heterodimers with RXR, a second nuclear receptor family member. PPARG/RXR heterodimers are activated when PPARG is bound by natural ligands (fatty acids and prostaglandins) or synthetic ligands including Troglitazone and Rosiglitazone. PPARG/RXR heterodimers binding to peroxisome proliferator response elements (PPREs), and without ligand, are maintained in an inactive state, in complexes with co-repressors (NCOR2, SMRT). Ligand binding induces a conformational change in the PPARG/RXR heterodimer causing the release of co-repressors and the recruitment of co-activators [(CREBBP, PPARGC1A and HAT), (Cohen, 2006; Nagy et al., 1999; Perissi et al., 1999)].

Studies in knockout mice indicate that Pparg regulates urothelial differentiation both in the ureter and bladder (Bell et al., 2011; Liu et al., 2019). In urothelial cell culture, Pparg agonists Troglitazone and Rosiglitazone in combination with an EGFR inhibitor, suppress squamous differentiation and induce expression of luminal markers, including *Upk1a*, *Upk2*, and *Krt20* (Varley and Southgate, 2008; Varley et al., 2004).

*PPARG* mutations and genomic alterations are common in bladder cancer. PPARG expression is downregulated in basal/squamous subtype tumors suggesting that loss of signaling may promote bladder tumor formation. To address this, we previously generated mice lacking Pparg throughout the urothelium using the *ShhCre* driver. These studies revealed a number of profound changes in the urothelium, including squamous metaplasia and loss of endogenous urothelial populations, likely due to alterations in the differentiation program of K14-Basal progenitors that produced squamous epithelial cells instead of urothelial cells. These observations suggest that Pparg is normally required for specification of K14-Basal cells, however, inactivation of *Pparg* alone during homeostasis is not sufficient to drive bladder cancer (Liu et al., 2019).

Activation of Pparg-dependent transcription either due to mutations in its binding partner RXR, or amplifications of the *PPARG* gene occur in 20-25% of luminal tumors (Halstead et al., 2017). These observations prompted us to examine whether gain-of-function mutations in Pparg could induce luminal subtype bladder cancer in mice. To do this, we inserted a cassette containing the HSV VP16 activator fused to the amino terminal of Pparg1 (Sugii et al., 2009) into the *Rosa26* locus where it is activatable in cells expressing Cre recombinase. We find that expression of *VP16;Pparg* in basal progenitors during homeostasis drives them to adopt a luminal (I-cell/S-cell) differentiation program *in situ*. These newly generated cells downregulate basal cell genes including *Krt5*, *Trp63*, and *Col17a1*, which mediates attachment to hemidesmosomes, after which they move to the upper layers of the urothelium and exit the cell cycle, but they don’t form tumors.

Exposure to carcinogens such as N-butyl-N-(4-hydroxybutyl)-nitrosamine (BBN), a nitrosamine found in tobacco smoke induces basal subtype lesions in mice after 4-5 months of exposure. BBN forms DNA adducts that result in DNA damage and mutations. BBN induces a potent inflammatory response during the first month of exposure that recedes in the following months as tumors develop (Vasconcelos-Nobrega et al., 2012). We find that the K14-basal progenitor population undergoes a damage response during the first month of BBN treatment in which they enter a transient activation state. This activation state is characterized by upregulation of defensins, including *Krt6a* and *Krt16*, which are not expressed in the healthy urothelium, as well as increased proliferation; similar to what is observed in the skin and airways in response to injury (Freedberg et al., 2001; Haensel et al., 2020; Zhang et al., 2019). Expression of the *VP16;Pparg* transgene in these activated K14-Basal cells results in a very different response compared to that in homeostasis: luminal tumors form after 4 months, while controls which do not express the *VP16;Pparg* transgene form basal/squamous tumors. These observations suggest that *Pparg* can drive activated basal cell progenitors to adopt a luminal differentiation program. Interestingly, we observed a shift at the base of some luminal lesions that increased with time, where luminal tumor cells downregulate luminal markers and begin to express basal markers, suggesting that tumors undergo a luminal to basal shift. Together our studies provide a model for studying luminal tumor formation *in vivo* and may also shed light on the mechanisms underlying tumor evolution, a phenomenon that has been reported in recent studies (Lamy et al., 2016; Lee et al., 2018).

## Results

### *Pparg* activation in basal cells induces a S-cell differentiation program

The *PPARG* gene is amplified in tumors of the Luminal subtype, which also express high levels of FABP4, a direct transcriptional target of *PPARG* (Choi et al., 2014). This suggests that *PPARG* is both over-expressed and transcriptionally active, most likely by endogenous ligands. To generate a gain-of-function model that mirrors the increased *PPARG* activity in luminal tumors, we generated mice harboring a constitutively active form of Pparg1 that is Tamoxifen inducible. We inserted a cassette containing the HSV *VP16* activator fused to the N-terminal of Pparg1 into the *Rosa26* locus where it is activatable in cells expressing Cre recombinase (Fig.1L). Unlike endogenous *Pparg*, *VP16;Pparg* is transcriptionally active without ligand binding (Saez et al., 2004; Sugii et al., 2009). K14-Basal cells have been shown to be progenitors that can produce tumors in mice (Papafotiou et al., 2016). To target this population, *VP16;Pparg* mice were crossed with the *Krt5Cre^ERT2^* line (hereafter referred to as *K5VP16;Pparg* mice), in which Tamoxifen-inducible Cre is expressed in K5-Basal cells and K14-Basal progenitors driven by the K5 promoter (Indra et al., 1999).

**Figure 1.**
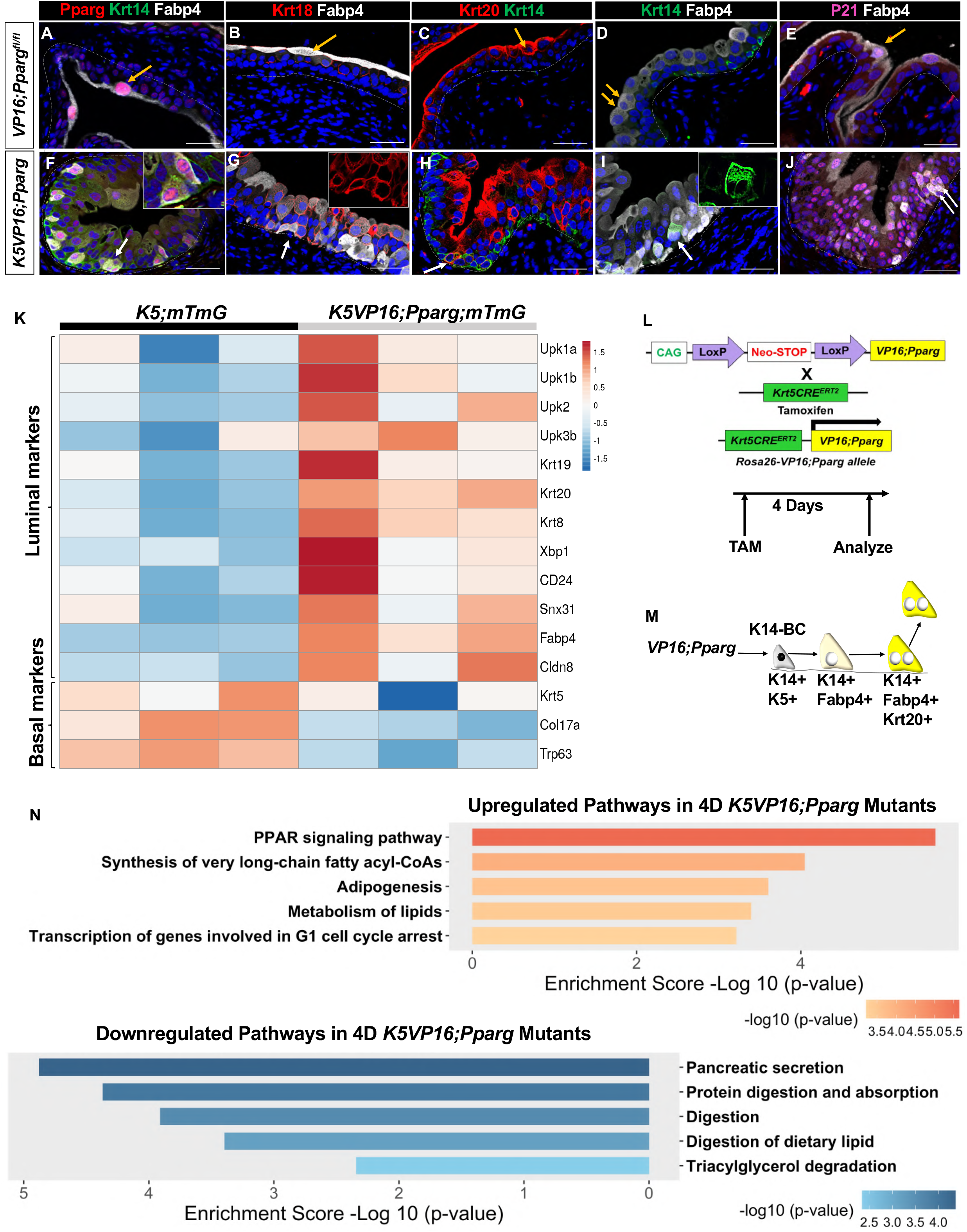
Expression of *VP16;Pparg* in basal cells induces an S-cell differentiation program. (A-J) Expressions of Pparg, Krt14, Fabp4 (A, F); Krt18, Fabp4 (B, G); Krt20, Krt14 (C, H); Krt14, Fabp4 (D, I); and P21, Fabp4 (E, J) in *VP16;Pparg* controls (A-E) and *K5VP16;Pparg* mutants (F-J). Yellow arrows denote Superficial cells in controls. White arrows denote mutant cells undergoing basal to luminal shift. The right-hand panels show higher magnifications of the areas marked by arrows. Scale bars, 50 μm. (K) Heatmap of luminal/basal gene signature in *K5;mTmG* controls and *K5VP16;Pparg;mTmG* mutants 4 days after Tamoxifen induction. (L) Schematic of mouse model and analysis. The *VP16;Pparg* mouse was crossed to basal Cre driver *Krt5Cre^ERT2^* in order to express the *VP16;Pparg* transgene in K5-basal cells. Bladders were harvested 4 days after the last Tamoxifen induction. (M) Schematic of S-cell differentiation program induced by VP16;Pparg expression in basal cells. (N) Upregulated and downregulated signaling pathways in *K5VP16;Pparg;mTmG* mutants compared to *K5;mTmG* controls 4 days after Tamoxifen induction.

Tamoxifen was administered intraperitoneally 3 times over the course of one week, which induced recombination in 80% of basal cells. To determine whether the VP16;Pparg mutant protein was transcriptionally active, we analyzed the distribution of Fabp4, a direct transcriptional target of *Pparg* in *K5VP16;Pparg* mutants and *VP16;Pparg^fl/fl^* controls 4 days after Tamoxifen induction. In controls Fabp4 and Pparg were co-expressed in S-cells as expected (Fig.1A), however in *K5VP16;Pparg* mutants, Fabp4 and Pparg were expressed both in S-cells in the luminal layer and in K14-basal cells in the basal layer (Fig.1F, Supplementary Fig.1R), indicating that the *VP16;Pparg* transgene is active. Further analysis revealed Krt18 and Krt20 which are selectively expressed in S-cells of controls (Fig.1B,C), co-localized with Fabp4 and Pparg in K14 positive cells in the basal layer (Fig.1G,H), suggesting that the *VP16;Pparg* transgene is inducing K14-Basal cells to differentiate into S-cells *in situ*. Consistent with this, we observed binucleated cells co-expressing K14 and Fabp4 in the basal layer, suggesting that this newly formed population has undergone failed cytokinesis, a process observed when I-cells differentiate into S-cells, which normally occurs in the luminal layers of the urothelium [Fig.1D,I, (Wang et al., 2018)].

Lineage tracing confirmed that K14+/Fabp4+/Krt20+ cells that formed in *K5VP16;Pparg* mutants were derived from basal cells. The *R26RmTmG* reporter (Muzumdar et al., 2007) was introduced into mutants and controls to generate *K5VP16;Pparg*;*mTmG* mice and *K5*;*mTmG* mice, respectively. In controls, Gfp expression was confined to the basal layer as expected (Supplementary Fig.1M,N,O,P), while in *K5VP16;Pparg* mutants, Gfp expressing cells which were observed in the basal layer 1 day after Tamoxifen induction, and Gfp expression expanded to encompass the entire urothelium by one 1 month (Supplementary Fig.1Q,R,S,T)

To identify changes in the transcriptional program of basal cells induced by expression of the *VP16;Pparg* transgene we performed bulk RNA-seq analysis on Gfp+ cells collected from *K5VP16;Pparg;mTmG* mutants and *VP16;Pparg^fl/fl^*;*mTmG* controls 4 days post Tamoxifen-induction. Pathway analysis revealed upregulation of *Pparg* signaling in basal cells of mutants, confirming that the transgene was active (Fig.1N). This analysis also revealed upregulation of S-cell markers in basal cells of mutants compared to controls, while *Krt5*, *Col17a* and *Trp63* which are highly expressed in basal cells of controls, were downregulated in mutants (Fig.1K), which was confirmed by immunostaining (Supplementary Fig.1U-X). These results suggest that that expression of the *VP16;Pparg* transgene in basal cells results in production of S-like cells *in situ,* in the basal layer of the urothelium. RNA-seq analysis also revealed changes in cell cycle genes including p21 (*Cdkn1a*), which was upregulated in basal cells from *K5VP16;Pparg* cells compared to *VP16;Pparg* controls and confirmed by immunostaining (Fig.1E,J), indicating that expression of the *VP16;Pparg* transgene induced S-cell like daughters to leave the cell cycle, as is the case with endogenous S-cells.

Squamous metaplasia in the airways, bladder and prostate is induced by Vitamin A-deficiency (Wilson et al., 1953; Wilson and Warkany, 1948). In this case, the urothelium is populated by cells expressing markers found in the skin, Krt13, Krt1, Krt10, similar to the phenotype induced in the urothelium in *Pparg* mutants, which is characterized by increase in K14-progenitors accompanied by a decrease in I-cells and S-cells (Liu et al., 2019). Retinoids have also been shown to be required for urothelial differentiation and regeneration, specifically for formation of I-cells and S-cells (Gandhi et al., 2013). We observed increased retinoid signaling in *K5VP16;Pparg* mutants compared to *VP16;Pparg*^fl/fl^ controls (Supplementary Fig.1Z). Upregulated genes include *Rbp4*, a lipocalin family member that transports retinol (the inactive form of Vitamin A) from the liver to tissues; *Stra6*, a receptor that transports retinol into the cell; *Rdh11* which synthesizes retinaldehyde from retinol in the first step of RA-synthesis; and *Aldh1a3*, a retinaldehyde dehydrogenase that converts retinaldehyde to RA, in the second step of RA synthesis [Supplementary Fig.1Z (Cunningham and Duester, 2015; Ghyselinck and Duester, 2019)]. These data suggest that *Pparg*-dependent RA-signaling is likely important for differentiation of luminal cell types, and for suppression of squamous differentiation.

Consistent with the known role of *Pparg* as a regulator of fatty acid transport and metabolism, we observed an increase in genes important in lipid metabolism (Fig.1N). These include *Scd1, Elovl, Acsl4, Cpt1, Fabp5, Fabp1*, *Lpl,* and *Cd36*. Genes involved in pancreatic secretion, protein digestion and absorption, digestion of dietary lipid, and triacylglycerol degradation were downregulated (Fig.1N). These pathways are similarly altered in non-alcoholic fatty liver disease [(NAFLD) (HM et al., 2020; Hoang et al., 1999; Lu et al., 2019; Suppli et al., 2019; Xiao et al., 2020)], which is thought to be linked to high fat diet induced *Pparg* signaling (Yu et al., 2003) Oil-Red-O staining of mutants 1 day, 4 days, and 1 month after Tamoxifen induction, revealed neutral triglyceride and lipid accumulation in mutant S-cells at 1 month, but staining was not detectable at earlier stages or in controls (Supplementary Fig.1A-L). These observations raise the possibility that the urothelium may respond to a high fat diet in a similar manner as observed in hepatocytes in steatosis, that turn on an adipocyte differentiation program in response to increased *Pparg* signaling.

Together these studies suggest that constitutive activity of *Pparg* in K14-Basal cell progenitors driven by *Krt5Cre^ERT2^* promoter induces formation of S-cell daughters instead of K5-Basal cells and I-cells, as normally occurs, a process that may depend on RA-signaling. With time, these S-cells appear to turn on an adipocyte differentiation program as occurs in NAFLD. These VP16;Pparg expressing S-cells are post-mitotic however, and hence did not form tumors.

### Short term treatment with carcinogens primes K14-Basal cells for tumor formation by inducing an activated state

Currently, the most established system for modeling MIBC in mice is N-butyl-N-(4-hydroxybutyl)-nitrosamine (BBN)(Cohen, 1998). BBN is a nitrosamine that is metabolized in the liver to N-butyl-N-(3-carboxypropyl)nitrosamine (BCPN), which is found in tobacco products (Bonfanti et al., 1988; Cohen, 1998; Mirvish, 1995). Five months of exposure to BBN induces basal subtype tumors, however short exposure induces a potent inflammatory response that resolves by about one month and recedes as tumors form (Degoricija et al., 2019). An interesting possibility is that this transient response to BBN might be important for priming urothelial cells, which are largely quiescent, to re-enter the cell cycle and produce tumors. To address this question, BBN was administered to wild type mice in water for 4 weeks, then we analyzed the urothelium from treated and untreated mice to determine the effects on urothelial populations as well as immune infiltration (Fig.2A). Histological analysis revealed edema and immune infiltration in the bladder of BBN treated mice which was not observed in controls, indicating the presence of inflammation (Fig.2B,C). Consistent with this, immunostaining revealed significant leukocyte infiltration in BBN-treated mice, as well as upregulation of p65 subunit of Nf-kB, a complex, that controls both innate and adaptive immunity (Fig.2D-G). These observations were confirmed by RNA-seq analysis, which revealed upregulation of T-cell markers (Cd4 and Cd8), proinflammatory cytokines (*Il1a*, *Il6*, *Il18*, *IFNg*, and *TNFa),* and transcription factors, including *Nf-kb, Foxp3*, *Stat3* and *Jun*, *Batf*, and *Fosl1,* which are *Ap1* family members that are important regulators of [(Atsaves et al., 2019), Fig.2P]

**Figure 2.**
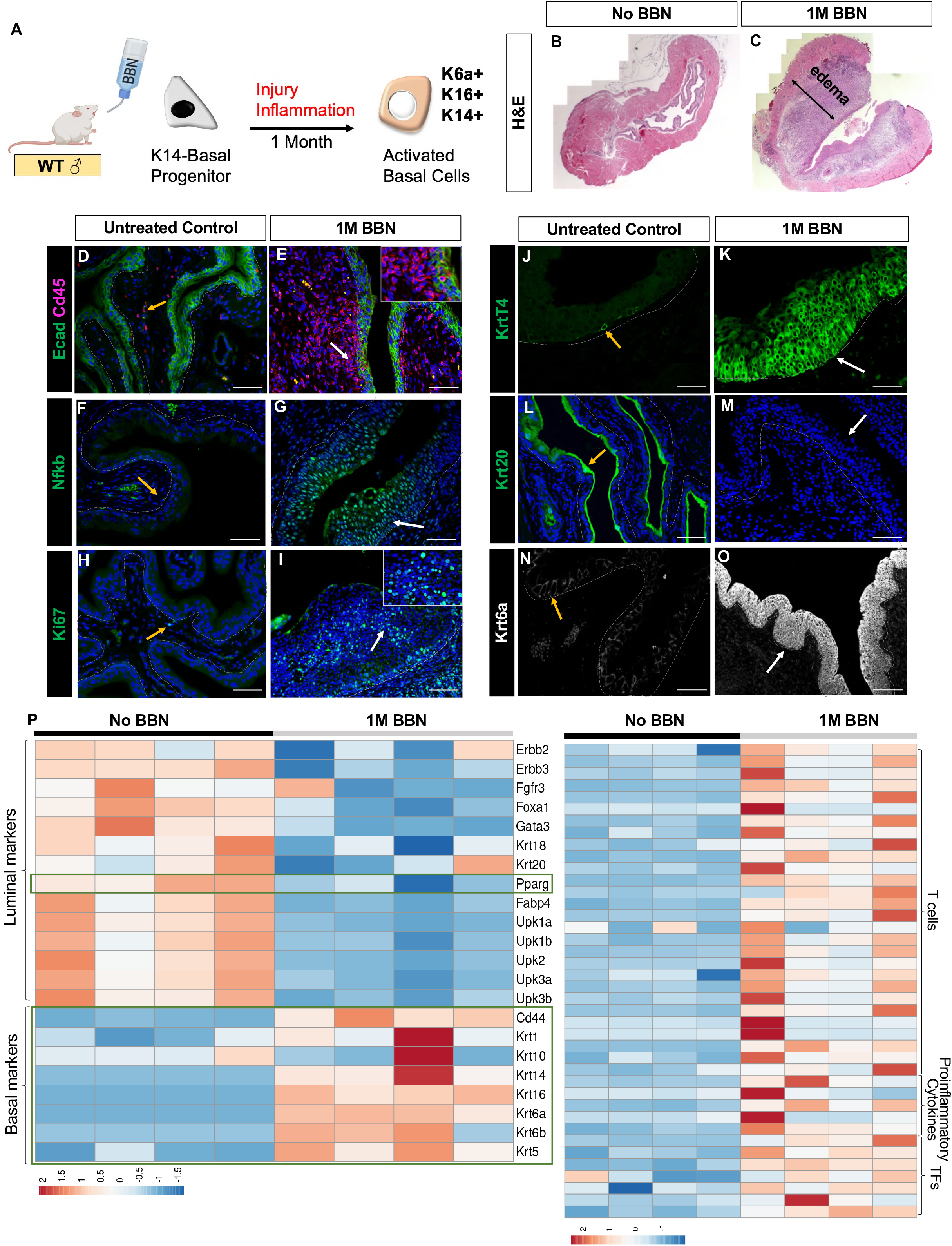
Short BBN treatment induces would healing response in the urothelium. (A) Schematic of activated basal cells after short BBN treatment. (B, C) H&E images of control (B) and 1 month BBN (C) bladders. Black double-headed arrow denotes edema in the stroma. (D-I) Expressions of E-cadherin, Cd45 (D, E); Nf-kb (F, G); and Ki67 (H, I) in control (D, F, H) and 1 month BBN (E, G, I) bladders. Yellow arrows denote expressions of Cd45 (D), Nf-kb (F), and Ki67 (H) before BBN treatment. White arrows denote expressions of Cd45 (E), Nf-kb (G), and Ki67 (I) after 1 month BBN treatment. The right-hand panels show higher magnifications of the areas marked by arrows. Scale bars, 50 μm. (J-O) Expressions of Krt14 (J, K), Krt20 (L, M), and Krt6a (N, O) in control (J, L, N) and 1 month BBN (K, M, O) bladders. Yellow arrows denote expressions of Krt14 (J), Krt20 (L), and Krt6A (N) before BBN treatment. White arrows denote expressions of Krt14 (K), Krt20 (M), and Krt6A (O) after 1 month BBN treatment. Scale bars, 50 μm. (P) Heatmaps of luminal/basal gene signatures and immune gene signatures in control and 1 month BBN bladders.

Further analysis by immunostaining and RNA-seq revealed widespread proliferation in the urothelium of BBN-treated compared to untreated controls, where proliferating cells were rare (Fig.2H,I). This short BBN-treatment also caused profound alterations in urothelial populations: K14-Basal cells which are rare in controls and are confined to the basal layer, now populated most of the urothelium in BBN treated mice (Fig. 2J,K). Consistent with the massive expansion of the K14-population, we observed depletion of I-cells and S-cells in BBN-treated mice, evidenced by downregulation of Krt20 (Fig. L,M) as well as *Fgfr3*, *Foxa1*, *Gata3*, *Krt18* and *Upks* (Fig.2P). On the other hand, BBN exposure induced upregulation of *Krt6a* which is barely detectable in the urothelium of healthy mice (Fig.2N,O) as well as keratins expressed in squamous epithelia including *Cd44*, *Krt1*, *Krt16*, *Krt6b* and *Krt5* (Fig.2P). K14-progenitors in the skin and airways, which are also barriers, upregulate *Krt14*, *Krt16* and *Krt6a* and undergo squamous differentiation. Hence, this activation state may be a conserved reaction to injury, perhaps maintaining the barrier by generating squamous tissue layer progenitors in response to injury.

### *VP16;PPARG* expression in activated basal cells induces luminal tumor formation

Long term BBN treatment of wild type mice results in basal subtype tumors (Fantini et al., 2018; Saito et al., 2018); however, a short BBN exposure activates the urothelium, which becomes proliferative and is populated almost exclusively with K14-Basal cells compared to untreated mice (Fig.2H-O). We tested whether expression of the *VP16;Pparg* protein after activation could alter the pathway of tumor formation from basal subtype to luminal subtype (Fig.3A). *K5VP16;Pparg* mutants and controls were exposed to BBN for 1 month, after which tamoxifen was administered intravesical under ultrasound guidance. Ultrasound analysis after 4 months revealed thickening of the bladder wall in controls and protrusions in the lumen of mutants (Fig.3B,C). H&E revealed papillary-like structures *K5VP16;Pparg* mutants and invasive lesions in *VP16;Pparg^fl/fl^* controls (Fig.3D,E). Pathological evaluation suggested that lesions in *K5VP16;Pparg* mutant mice were consistent with human Ta and T1 (papillary exophytic) lesions (Fig.3H,L,X) while control bladders without the *VP16;Pparg* transgene contained invasive T3 (squamous) lesions (Fig. 3P,T). Laminin and smooth muscle actin (SMA) staining allowed us to clearly visualize the fibrovascular core of luminal lesions, a hallmark of papillary (luminal) lesions while in controls, vasculature was widespread (Fig.3F,G). To confirm that lesions derived from basal cells in mutants and controls, we analyzed bladders from *K5VP16;Pparg;mTmG* mutants and *K5;mTmG* controls 4 months after tamoxifen induction, when lesions were clearly visible. In both cases, lesions were almost completely Gfp-positive (Supplementary Fig.3A,B), indicating that they are derived from Gfp+ basal cells.

**Figure 3.**
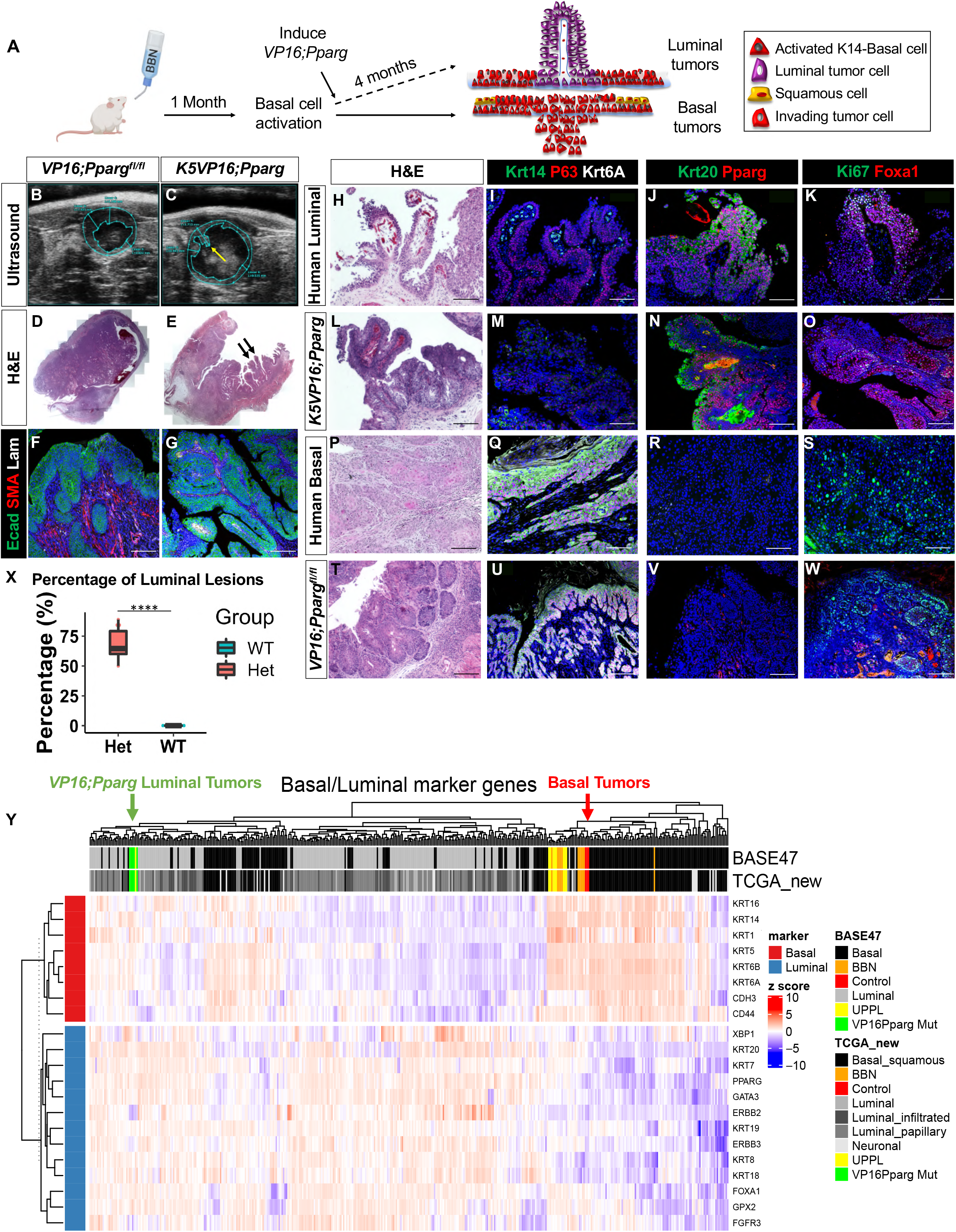
Activation of *Pparg* in *K5VP16;Pparg* mice produces luminal bladder tumors. (A) Schematic of activating *Pparg* in *K5VP16;Pparg* mice to produce luminal tumors. (B-G) Ultrasound images of bladders 4 months after Tamoxifen induction in *VP16;Pparg^fl/fl^* control (B) and in *K5VP16;Pparg* mutant (C). Yellow arrow denotes lesion protruding into the lumen. H&E images (D, E) of bladders 4 months after Tamoxifen induction in *VP16;Pparg^fl/fl^* control (D) and in *K5VP16;Pparg* mutant (E). Black arrows denote high grade papillary lesions. Expressions of E-cadherin, smooth muscle actin, and Laminin (F, G) in *VP16;Pparg^fl/fl^* control (F) and in *K5VP16;Pparg* mutant (G). Scale bars, 100 μm. (H-W) H&E images of human luminal tumor (H), *K5VP16;Pparg* mutant tumor (L), human basal tumor (P), and *VP16;Pparg* control basal tumor (T). Expressions of Krt14, P63, Krt6Aa(I, M, Q, U); Krt20, Pparg (J, N, R, V); Ki67, Foxa1 (K, O, S, W) in human luminal tumor (I-K), *K5VP16;Pparg* mutant tumor (M-O), human basal tumor (Q-S), and *VP16;Pparg* control basal tumor (U-W). Scale bars, 100 μm. (X) Quantification of percentage of luminal tumors observed in *VP16;Pparg* controls (n=10) and *K5VP16;Pparg* mutants (n=10). Significance calculated by Mann-Whitney U test. ****p≤0.0001 (Y) Heatmap of *K5;mTmG* control tumors, *K5VP16;Pparg;mTmG* mutant tumors, BBN, and UPPL primary tumor samples with the TCGA BLCA dataset across luminal/basal genes using the BASE47 classifier. Green arrow denotes *VP16;Pparg* luminal mutant tumors. Red arrow denotes control basal tumors.

Comparison of BBN-induced lesions in *K5VP16:Pparg* luminal lesions with human luminal tumors from patients revealed similar branched exophytic structure and a fibrovascular core in both. Conversely, BBN-treated control mice lesions were invading into the submucosa and muscle, similar to the human basal tumor (Fig 3H,L,P,T). Immunostaining of human and mouse lesions with markers expressed in basal subtype lesions (Krt14 and Krt6a) revealed robust expression in the BBN-treated controls and human basal tumor, whereas there was little if any expression in lesions from mouse *K5VP16;Pparg* mutants or luminal lesions from human patients (Fig.3I,M,Q,U). Analysis with markers expressed in luminal subtype tumors (Krt20, Pparg, Foxa1) revealed little expression in control BBN-induced basal lesions and human basal tumors, while expression was prominent in luminal human tumors and *K5VP16;Pparg* tumors (Fig.3J,K,N,O,R,S,V,W). Expression of Ki67, which marks proliferating cells, was distinct in basal and luminal tumors. In basal tumor, Ki67 expression was in cells adjacent to the basement membrane of lesions, while in luminal tumors proliferating cells were scattered (Fig.3K,O,S,W).

RNA-seq analysis of laser-captured cells from *K5VP16;Pparg* and control mice revealed co-clustering with genes expressed in tumors with luminal and basal subtypes, respectively. Comparison of the transcriptomes of BBN-induced tumors with subtypes from Cancer Genome Atlas (TCGA) human tumors (n=408) and known luminal bladder cancer model *Upk3a-Cre^ERT2^; Trp53L/L; PtenL/L; Rosa26LSL-Luc* (UPPL) tumors and controls reveal that the *K5VP16;Pparg* mutant lesions co-clustered with Luminal and Luminal Papillary TCGA samples, which express a set of markers including *Pparg, Foxa1, Krt18, and Krt20* (Fig.3Y). Control basal lesions from BBN-treated *VP16;Pparg^fl/fl^* mice co-clustered with the Basal Squamous TCGA samples, which express a set of markers including *CD44, Krt5, Krt14,* and *Krt6a* (Fig.3Y). Pathway analysis revealed upregulation of genes related to metabolism in *K5VP16;Pparg* mutant lesions (Supplementary Fig.3E), which is not surprising given the known role of Pparg as a regulator of mitochondrial biogenesis and fatty acid metabolism. Interestingly, *Foxa1* and *Fgfr3* pathways are upregulated in luminal tumors from *K5VP16;Pparg* mice (not shown). Both *Foxa1* and *Fgfr3* have been shown to be upregulated in the luminal subtype of bladder cancer (DeGraff et al., 2012). In particular, FOXA1 is associated with high grade, late-stage bladder cancer and increased tumor proliferation (Osei-Amponsa et al., 2020). Pathways related to cell cycle, T-cell activation, and cytokine signaling are downregulated in the *K5VP16;Pparg* mutant lesions, suggesting the immune response is dampened in the *Pparg* activated luminal tumors (Supplementary Fig.3E). Taken together, these findings suggest activating *Pparg* in activated basal cells can induce basal tumors to adopt a luminal fate.

### *Pparg* induced luminal tumors are immune excluded

An important distinction in the classification of tumors is whether the tumor is immune “hot” or immune “cold.” Hot tumors are characterized by increased immune cell trafficking, an abundance of inflammatory cytokines and antigen presenting cells, increased T-cell activation, and increased major histocompatibility complex (MHC 1) expression. Cold tumors have low to no immune cell trafficking, impaired T-cell activation, an abundance of myeloid derived suppressor cells (MDSCs), and regulator T-cells which release immunosuppressive cytokines (Noman et al., 2020; Rijnders et al., 2017). Human basal and luminal MIBCs demonstrate varying degrees of immune infiltration. Immune infiltration is mostly associated within the basal/squamous and stroma-rich subtypes of MIBC, whereas the luminal subtypes are often immune excluded (Kamoun et al., 2020), suggesting that the tumor microenvironment in bladder cancer is driven by subtype-specific differences. However, whether the immune exclusion phenotype is due to active immune suppression in the luminal subtypes is largely unknown. *Pparg* is known to regulate a number of immune responses, including Nf-kb suppression and interactions with the AP1 pathways (Chung et al., 2000; Korpal et al., 2017; Remels et al., 2009; Ricote and Glass, 2007; Ricote et al., 1998). Unsupervised clustering of *K5VP16;Pparg* mutants and *VP16;Pparg^fl/fl^* control tumors using previously established immune gene signature demonstrated overall low levels of immune response, similar to the UPPL luminal bladder cancer model [(Fig.4A, (Saito et al., 2018)]. In contrast, the control tumors showed very active immune signatures (Fig. 4A). To further investigate these findings, we compared expressions of Nf-kb subunit p65 and Cd45 in *K5VP16;Pparg* mice and *VP16;Pparg^fl/fl^* controls (Fig.4B-E). Immunostaining of control basal tumors revealed widespread nuclear expression of p65, indicating that Nf-kb is activated throughout the tumor. Additionally, we observed numerous infiltrating Cd45+leukocytes (Fig.4C,E). Analysis of p65 expression in *K5VP16;Pparg* mutant tumors, reveal little if any nuclear expression, suggesting that Nf-kb is inactive in the mutants. Likewise, we did not observe infiltrating immune cells in these lesions based on Cd45 expression. *Pparg* actively represses Nf-kb expression suggesting that Pparg is likely to be important for inducing the immune excluded phenotype in luminal tumors.

**Figure 4.**
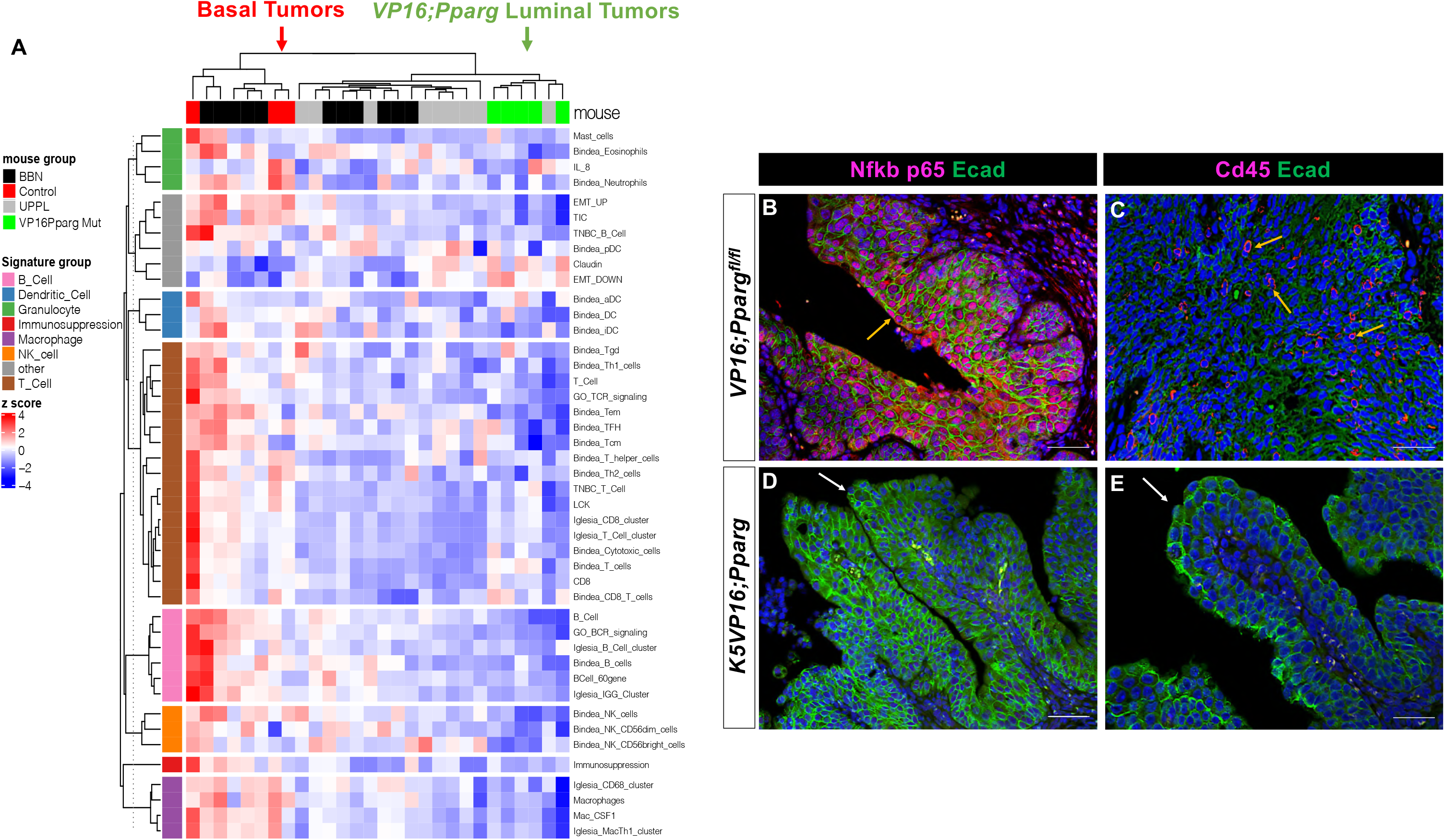
*K5VP16;Pparg* tumors are immune excluded. (A) Immune gene signatures of *K5;mTmG* control tumors, *K5VP16;Pparg;mTmG* mutant tumors, BBN, and UPPL primary tumor samples. Green arrow denotes *VP16;Pparg* luminal mutant tumors. Red arrow denotes control basal tumors. (B-E) Expressions of Nf-kb p65, E-cadherin (B, D); and Cd45, E-cadherin (C, E) in *VP16;Pparg* control tumors (B, C) and *K5VP16;Pparg* mutant tumors (D, E). Yellow arrows denote control tumor cells with Nf-kb p65 expression (B) and Cd45 lymphocyte infiltration (C). White arrows denote the absence of Nf-kb p65 expression (D) and Cd45 lymphocyte infiltration (E) in *K5VP16;Pparg* mutant tumors. Scale bars, 100 μm.

### A subset of Luminal tumors in *K5VP16;Pparg* mutants undergo a shift toward the basal subtype

During the course of our analysis, we observed a domain in a few luminal tumors that appears to be more basal in character than luminal (Fig.5A,F). Mixed subtypes were observed n 15/70 lesions in *K5VP16;Pparg* mutants 4 months after Tamoxifen induction (Table1;Fig.5A,F). However, these mixed subtype tumors were rare in animals analyzed 1 month earlier, where only 1/28 lesions displayed a mixed subtype (Table 1), suggesting that the shift increases with time. Analysis with Krt6a and Krt14, markers that stain basal subtype tumors, revealed robust expression throughout basal lesions in controls, which were exposed to BBN for 5 months but do not express the *VP16;Pparg* transgene (Fig.5B). Analysis of Krt6 and K14 expression in tumors with mixed histology revealed little staining in the luminal compartment (Fig.5G, green bracket), while staining was increased in the basal-like compartment (Fig.5G, blue bracket). Foxa1, a luminal marker was not detectable in control basal subtype tumors (Fig.5C), however expression was robust in the luminal portion of the mixed subtype tumor (Fig.5H, green bracket). Foxa1 expression was maintained in the basal-like portion of this tumor, but expression was lower than in the luminal portion (Fig.5H, green and blue bracket denote the luminal and basal-like portions of the tumor). Similar findings were observed with Pparg; expression was undetectable in basal control tumors (Fig.5D,E), while expression was maintained at high levels in the luminal upper domain (Fig.5I,J, green bracket). In the basal-like portion of these mixed lesions, however, levels were lower compared to the luminal domain (Fig.5I,J blue bracket, inset shows a higher magnification of *Pparg* expression in the basal-like domain).

**Figure 5.**
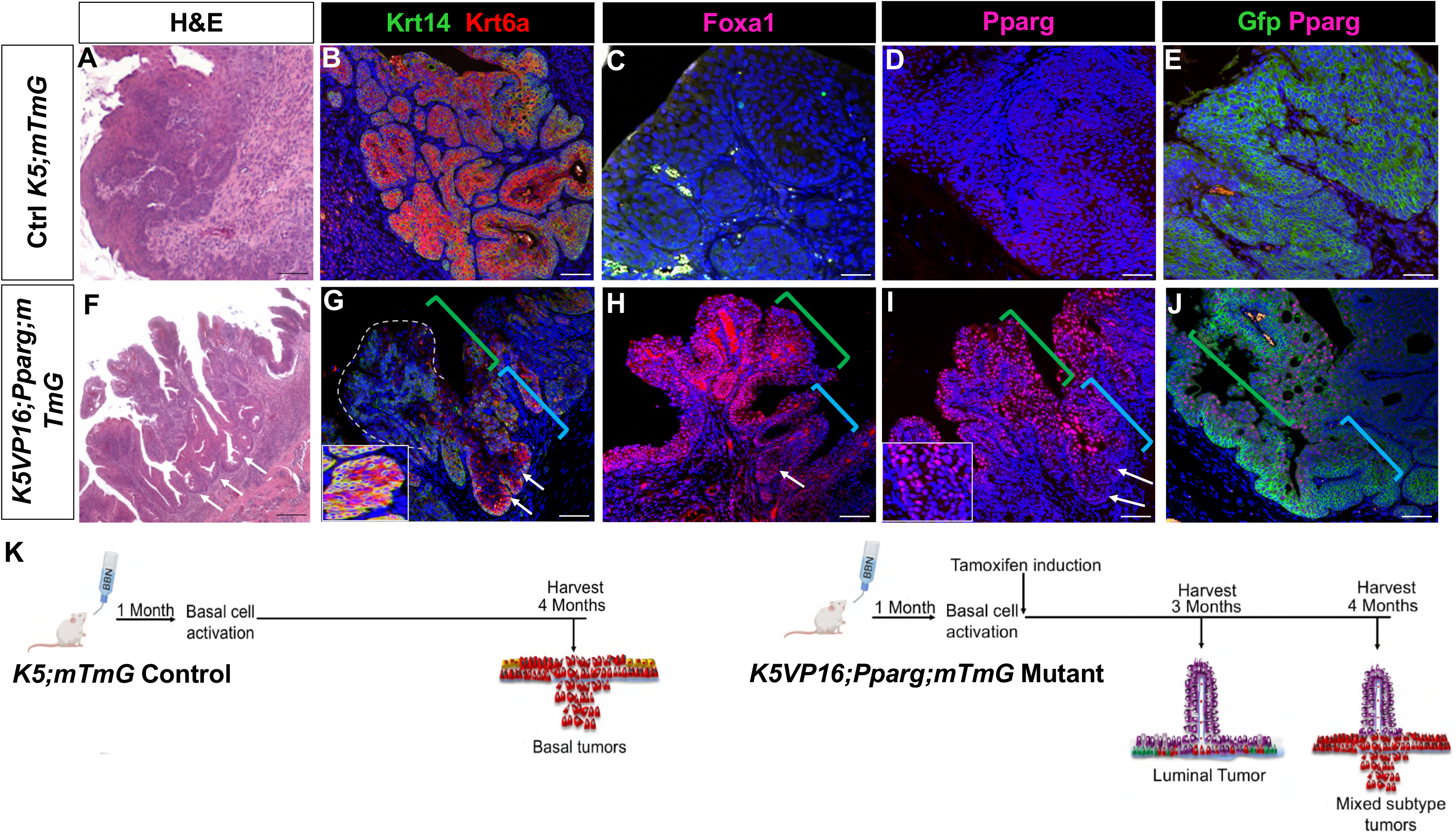
*K5VP16;Pparg* tumors have luminal and basal domains. (A-J) H&E images of *K5;mTmG* control tumor (A) and *K5VP16;Pparg;mTmG* mutant (F) tumor at 4 months. Expressions of Krt14, Krt6A (B, G); Foxa1 (C, H); Pparg (D, I); and Gfp, Pparg (E, J) in *VP16;Pparg* control tumors (B-E) and *K5VP16;Pparg* mutant tumors (G-J) at 4 months. Green brackets indicates mutant luminal domain at 4 months. Blue bracket indicates mutant basal domain at 4 months (D). White arrows denote mutant basal domain at 4 months. The left-hand panel in (G) shows a neighboring invasive basal lesion from the same animal. The left-hand panel in (I) shows higher magnifications of the areas marked by arrows. Scale bars, 100 μm. (K) Schematic of tumor evolution in *K5;mTmG* controls and *K5VP16; Pparg;mTmG* mutants.

**Table 1.** Quantification of tumor class in *VP16;Pparg* controls and *K5VP16;Pparg* mutants. Overall classification and quantification of lesions observed in *VP16;Pparg* controls and *K5VP16;Pparg* mutants at the indicated timepoints. Pathologic classification is based on analysis of the H&E staining.

| Time point | Sample | # of Luminal Lesions |  | # of Basal Lesions | Time point | Sample | # of Luminal Lesions |  | # of Basal Lesions |
| --- | --- | --- | --- | --- | --- | --- | --- | --- | --- |
|  |  | Luminal only | Luminal with basal bottom |  |  |  | Luminal only | Luminal with basal bottom |  |
| **4 Months**<br>*K5VP16;Pparg* | Het 72 | 5 | 1 | 2 | **4 Months**<br>**Control** | WT 5 | 0 | 0 | 2 |
|  | Het 74 | 8 | 1 | 1 |  | WT 3 | 0 | 0 | 5 |
|  | Het 51 | 6 | 3 | 6 |  | WT 88 | 0 | 0 | 1 |
|  | Het 1 | 18 | 4 | 4 |  | WT 54 | 0 | 0 | 3 |
|  | Het 53 | 2 | 4 | 3 |  | WT 55 | 0 | 0 | 1 |
|  | Het 56 | 1 | 0 | 1 |  | WT 2 | 0 | 0 | 3 |
|  | Het 57 | 3 | 0 | 3 |  | WT 83 | 0 | 0 | 5 |
|  | Het 70 | 5 | 1 | 4 |  | WT 20 | 0 | 0 | 2 |
|  | Het 9 | 1 | 0 | 2 |  | WT 19 | 0 | 0 | 1 |
|  | Het 75 | 6 | 1 | 2 |  | WT 56 | 0 | 0 | 2 |
|  | **Total** | **55** | **15** | **28** |  | **Total** | **0** | **0** | **25** |
| **3 Months**<br>*K5VP16;Pparg* | Het 44 | 8 | 1 | 3 | **3 Months**<br>**Control** | WT61 | 0 | 0 | 3 |
|  | Het 62 | 6 | 0 | 3 |  | WT58 | 0 | 0 | 4 |
|  | Het 50 | 2 | 0 | 0 |  | WT59 | 0 | 0 | 4 |
|  | Het 4 | 7 | 0 | 1 |  | WT5 | 0 | 0 | 4 |
|  | Het 7 | 4 | 0 | 3 |  | WT81 | 0 | 0 | 3 |
|  |  |  |  |  |  | WT22 | 0 | 0 | 2 |
|  | **Total** | **27** | **1** | **10** |  | **Total** | **0** | **0** | **20** |
| **2 Months**<br>*K5VP16;Pparg* | Het70 | 4 | 0 | 0 | **2 Months**<br>**Control** | WT50 | 0 | 0 | 1 |
|  | Het56 | 1 | 0 | 0 |  | WT49 | 0 | 0 | 1 |
|  | Het62 | 2 | 0 | 0 |  | WT40 | 0 | 0 | 2 |
|  | **Total** | **7** | **0** | **0** |  | **Total** | **0** | **0** | **4** |
| **1 Month**<br>*K5VP16;Pparg* | Het 8 | 0 | 0 | 0 | **1 Month**<br>**Control** | WT 66 | 0 | 0 | 0 |
|  | Het 50 | 0 | 0 | 0 |  | WT 70 | 0 | 0 | 0 |
|  | Het 10 | 0 | 0 | 0 |  | WT 79 | 0 | 0 | 0 |
|  | **Total** | **0** | **0** | **0** |  | **Total** | **0** | **0** | **28** |

Lineage tracing of *K5;mTmG* controls, revealed *Gfp* expression throughout basal subtype tumors (Fig.5E). Analysis of *K5VP16;Pparg*;*mTmG* mice revealed Gfp-labeling both in the luminal domain and in the basal-like domain of mixed subtype tumors, indicating that both luminal and basal-like compartments arose from cells expressing the *K5VP16;Pparg*;*mTmG* transgene (Fig.5J). The observations that (i) the number of tumors with the mixed basal/luminal subtype increases with time, (ii) that these tumors express *Ppar*g both in luminal and basal-like domains, (iii) that both luminal and basal domains are Gfp-positive, suggests that lower basal-like domain is derived from the upper luminal tumor, and had begun to shift from a luminal to a basal subtype.

To further characterize changes in the mixed subtype lesions, we performed RNAseq analysis using laser capture to collect tissue from the luminal and basal-like domains separately, as well as from basal subtype controls. Analysis of basal subtype controls revealed high levels of expression of basal subtype markers, including *Cd44*, *Krt6a*, *Krt14* and *Krt16*, while expression of luminal markers was low (Fig.6A,D). Analysis of the upper luminal domain of mixed subtype lesions revealed a pattern of expression typical of luminal tumors; Basal markers were downregulated and luminal markers including *Pparg*, *Upks*, *Krt20*, *Gata3* and *Foxa1* were upregulated (Fig.6B,D). Expression of most luminal markers was low in the basal-like portion of the tumor compared to the luminal upper portion (Fig.6B,C) while a subset of basal markers were upregulated, consistent with the results of immunostaining (Fig.6C). Pathway analysis revealed increases in genes important for metabolism and lipid synthesis, as well as glutathione-mediated detoxification, a likely response to BBN exposure and the Pparg signaling pathway. Genes upregulated in the basal portion of the mixed subtype tumors include those important for cell cycle, T-cell signaling, and inflammation, which is consistent with the immune infiltrated phenotype of basal subtype tumors (Fig.6E). Taken together, our observations suggest that advanced luminal tumors lose Pparg expression, display increased expression of basal markers, and develop invasive basal domains. This shift in phenotype has recently been reported in patient-derived organoids (Lee et al., 2018) as well as in human tumors (Lamy et al., 2016). We have established a mouse model that undergoes a phenotypic shift from a luminal subtype towards a basal subtype. Luminal tumors are often classified as NMIBC, however muscle invasive lesions develop in 10-20% of patients diagnosed with luminal NMIBC (Kamoun et al., 2020; Knowles and Hurst, 2015; Tan et al., 2019). An important question is whether human invasive lesions develop from the original luminal tumor in a similar way as observed here, since luminal tumors are generally removed by TURBT, which may leave behind the basal-like invasive portion of the tumor.

**Figure 6.**
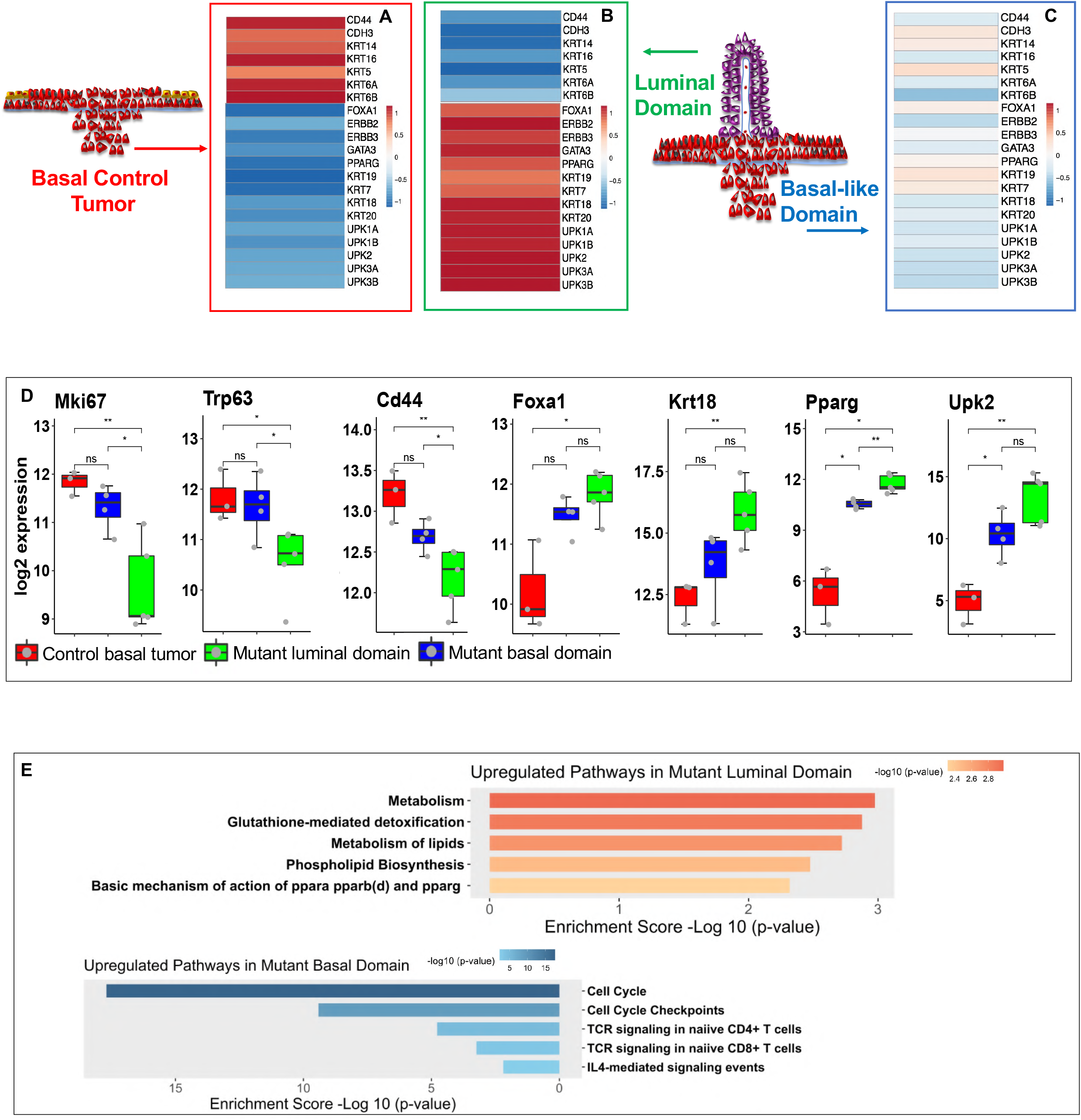
Evolution of *K5VP16;Pparg* tumors. (A-C) Heatmaps of luminal/basal gene signatures in *K5;mTmG* basal control tumors (A) and luminal (B) and basal (C) domains of *K5VP16;Pparg;mTmG* mutant tumors at 4 months. Red arrow and box indicate basal control tumor. Green arrow and box indicate mutant luminal domain. Blue arrow and box indicate mutant basal domain. (D) mRNA expression of *Mki67, Trp63, Cd44, Foxa1, Krt18, Pparg,* and *Upk2* from *K5;mTmG* basal control tumors, luminal, and basal domains of *K5VP16;Pparg;mTmG* mutant tumors at 4 months. Significance calculated by Mann-Whitney U test. ns=not significant; *p≤0.05; **0.05 ≤p≤0.01. (E) Upregulated and downregulated signaling pathways in *K5VP16;Pparg* mutant luminal domain compared to basal domain at 4 months.

## Discussion

*Pparg* has the capacity to regulate numerous cellular functions, including metabolism, fatty acid transport, and cell type specification. *PPARGPPARG* expression is downregulated in the basal subtype of bladder cancer, and amplifications of the *PPARG* gene or activating mutations in *R*XR, the PPARG binding partner, are present in about 17% of luminal tumors (Biton et al., 2014; Cancer Genome Atlas Research, 2014; Rochel et al., 2019), which has led to the suggestion that *PPARG* drives luminal tumor formation. The healthy urothelium is populated by a superficial layer of S-cells, which are post-mitotic, layers of I-cells that can replace S-cells when they die off during homeostasis or acute infection, K5-Basal cells which reside in the basal and suprabasal layers, and K14-Basal cells which are rare and are confined to the basal layer. Based on lineage studies with the *Krt5Cre^ERT2^* driver, K14-basal cells have the capacity to regenerate the urothelium *de novo* (Schafer et al., 2017b; Shin et al., 2011; Shin et al., 2014), and lineage studies together with a BBN model of carcinogenesis indicate that K14-Basal cells also give rise to bladder cancers (Papafotiou et al., 2016). Our studies indicate that Pparg plays an important role in specifying the differentiation program of K14-Basal cells. Our previous studies suggest that loss of Pparg results in impaired differentiation of I-Cells and S-cells, increased proliferation, and squamous metaplasia. In this case, the K14-Basal population expands and produces squamous cell types instead of endogenous urothelial populations (Liu et al., 2019). The studies described here indicate that active Pparg signaling can induce K14-Basal cells to differentiate into S-cells *in situ* and can also shift the differentiation program in basal subtype bladder cancer toward a luminal subtype. We also observe changes in the differentiation state of luminal tumors over time that suggest a luminal to basal shift is occurring. This shift is accompanied by downregulation of *Pparg* as well as other luminal markers. Whether downregulation of *Pparg* is the critical factor that induces these changes is an important question.

### Basal cell activation may be a hallmark of cancer in epithelial barriers

Cigarette smoking is a major risk factor for bladder cancer (Society, 2020). Several studies indicate that carcinogens in tobacco smoke induce formation of DNA adducts that lead to DNA damage and point mutations (Pfeifer et al., 2002; Weng et al., 2018). Point mutations can lead to production of neoantigens that are recognized as foreign by resident dendritic cells or macrophages triggering an inflammatory response. Expression of the *VP16;Pparg* transgene in K14-Basal cells during homeostasis induces a terminal differentiation program, however we find that a short period of exposure to BBN, a carcinogen or cyclophosphamide, which contains acrolein alters the urothelial microenvironment, inducing an inflammatory response that may prime K14-Basal cells for tumor formation.

Studies in the skin and airways indicate that K14-Basal cells, which are progenitors in both tissues, respond to injury or inflammation by taking on an activated state (Freedberg et al., 2001; Haensel et al., 2020; Shaykhiev, 2015). These cells produce squamous epithelial cells instead of epidermal cell types, which upregulate Krt6a and Krt16. We find a similar situation exists in the urothelium in response to a short period of carcinogen exposure. K14-Basal progenitors normally self-renew and produce K5-Basal cells that populate the basal and suprabasal layers, and I-cell daughters that differentiate into S-cells.We find that 1 month of BBN exposure or repeated exposure to cyclophosphamide induces an activation state in K14-Basal cells similar to that observed in the epidermis and airways. The K14-Basal population becomes proliferative, expands and ceases producing endogenous urothelial cell types (K5-Basal cells and I-cells), instead producing squamous cell daughters that express *Krt6a and Krt16*, which are not detected in the healthy urothelium.

The activation process in the skin is initiated by secretion of IL-1 and TNFa (Groves et al., 1995; Kondo and Ohshima, 1996), and is thought to be sustained by TNFa and TGF signaling, leading to activation of the NF-KB signaling pathway and the associated inflammatory responses (Ghosh and Karin, 2002; Lawrence, 2009; Solt et al., 2009; Solt et al., 2007). Consistent with this, we also observe upregulation of Il-1, Tnf, Ifng, and Nf-kb signaling in the bladder after short term BBN treatment. Inflammatory reactions have long been implicated as a precursor prior to the formation of tumors in several types of cancer (Candido and Hagemann, 2013), which may be a response to neoantigen production. Therefore, the identification of activated basal cells as the cell population that responds to inflammation suggests that entry to the activated basal cell cycle is a necessary step for tumor formation.

### Potential mechanisms of Immune evasion in VP16;Pparg induced luminal tumors

RNA-seq analysis reveals lack of upregulation of a large number of immune mediators in luminal tumors in *K5VP16;Pparg* mutants and are immune excluded. In some types of cancer, with immune suppression, immune cells are observed near tumors, but fail to penetrate. In luminal tumors of *K5VP16;Pparg* mutants however, immunostaining reveals few if any immune cells in or near tumors, while control basal subtype tumors are immune infiltrated.

Nf-kb is a master regulator of the immune response, a direct Pparg target that is actively suppressed by Pparg signaling (Remels et al., 2009; Scirpo et al., 2015). We find that Nf-kb (p65 expression), which is abundant in basal subtype tumors is barely detectable in luminal tumor cells, suggesting that suppression of Nf-kb may be an important factor in the immune excluded phenotype.

### Tumor evolution

Intra-tumor heterogeneity and evolution have been observed in bladder cancer studies of tumor-derived organoids and mutational analysis of human tumors (Heide et al., 2019; Lee et al., 2018; Warrick et al., 2019). We observe altered histology at the base of luminal tumors in *K5VP16;Pparg* mutants that appears 4 months after Tamoxifen induction, but not before, suggesting that this phenotype is acquired over time. These domains express basal markers including Krt6a, and downregulate luminal markers. It is likely that new mutation acquisition from sustained exposure to BBN after Tamoxifen induced expression of the *VP16;Pparg* transgene may be a driver of these changes, as bladder cancer has one of the highest tumor mutational burden. However, the observation that they form, and are in a basal position suggests that they may not be observed in biopsies of luminal subtype tumors taken from patients, or may remain after TURBT treatment, thus contributing to recurrence and progression from NMIBC to MIBC.

**Key Resources Table.**
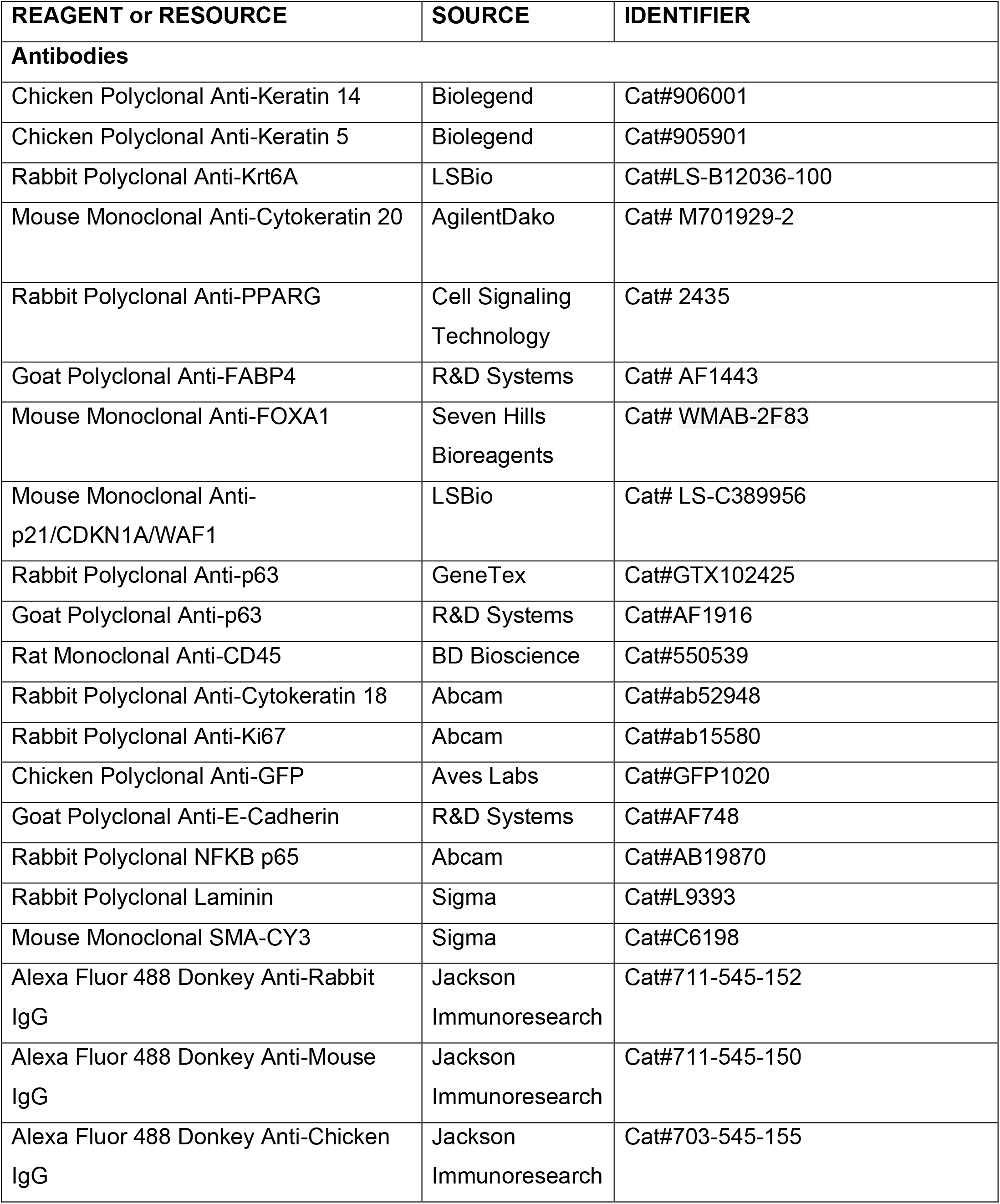
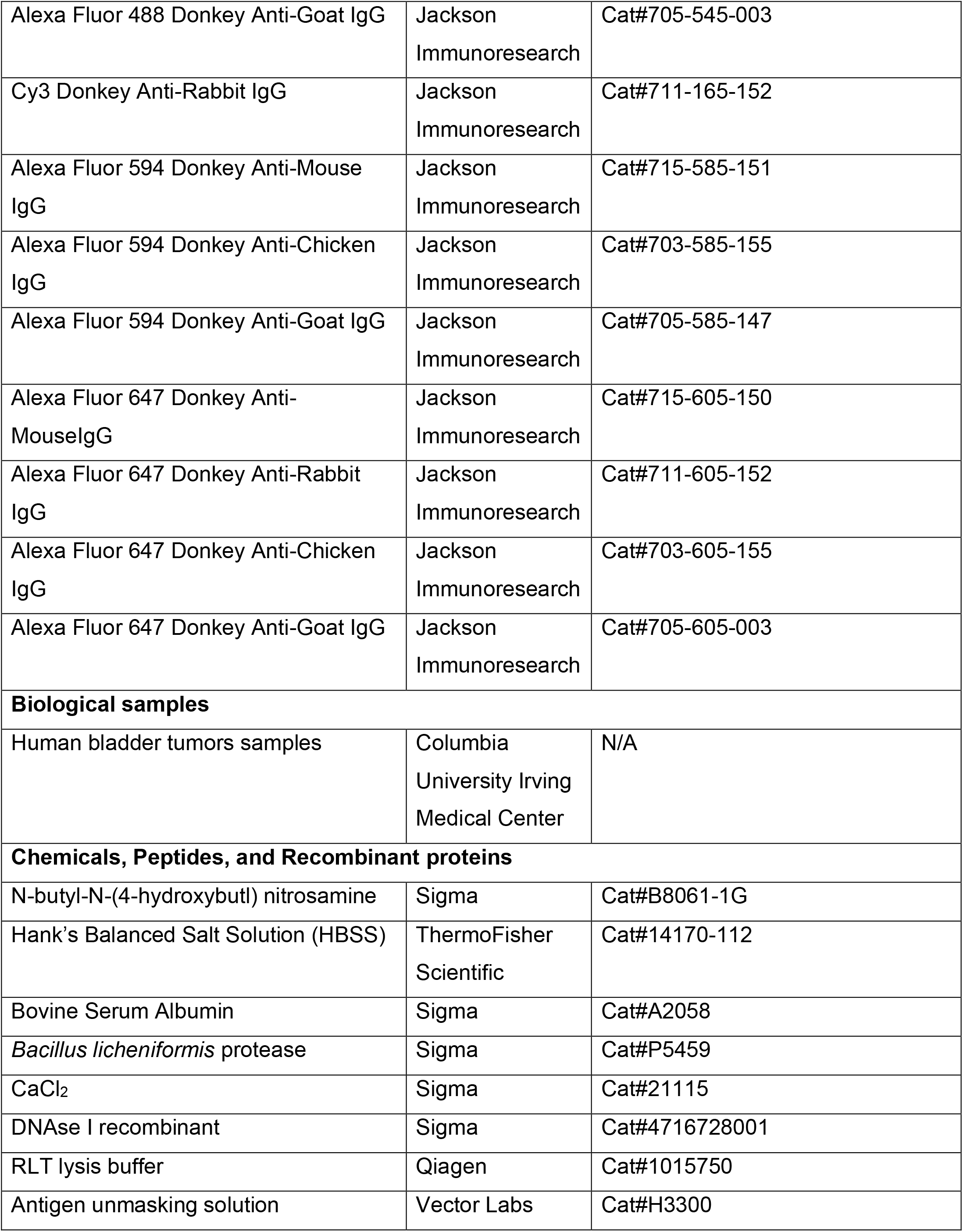
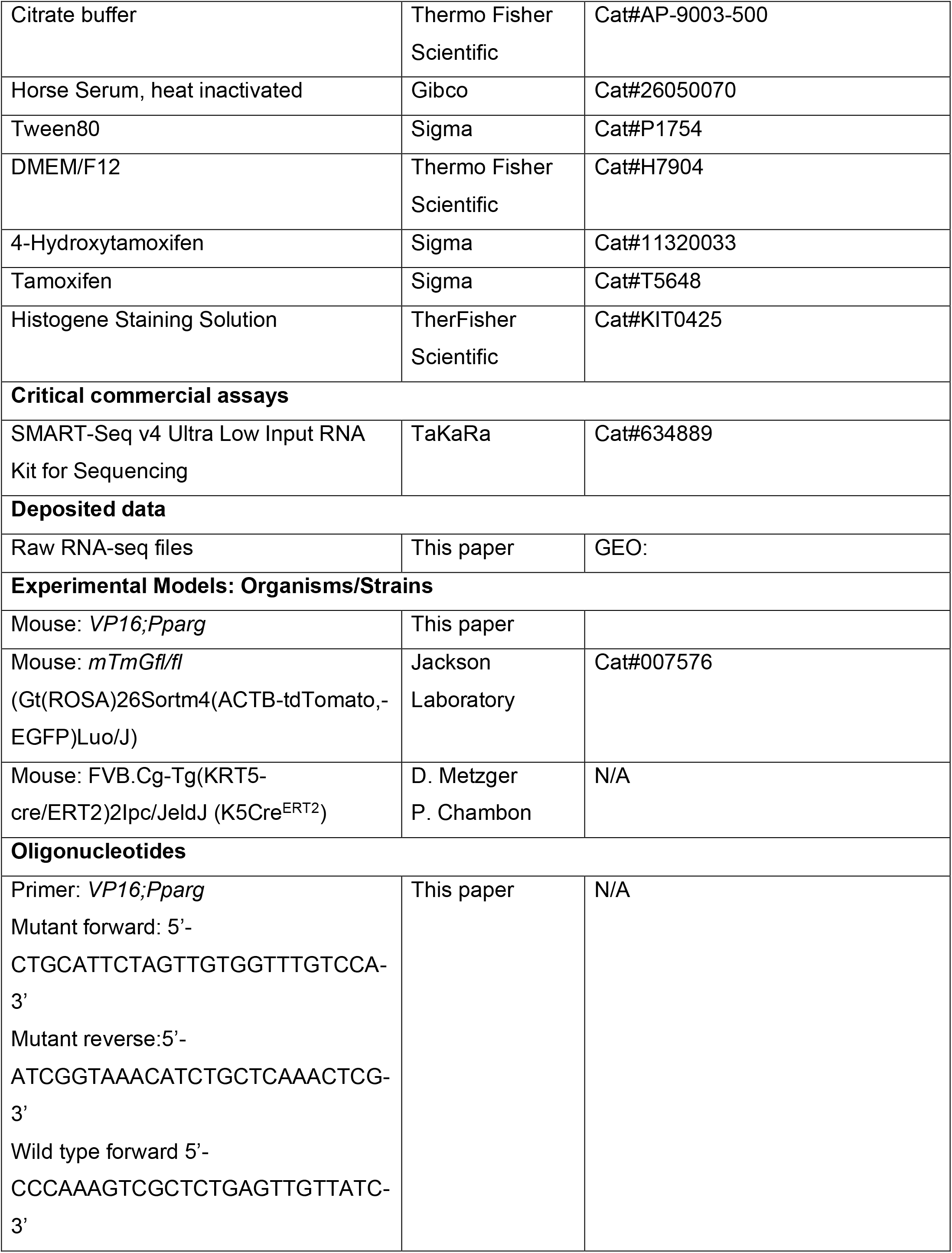
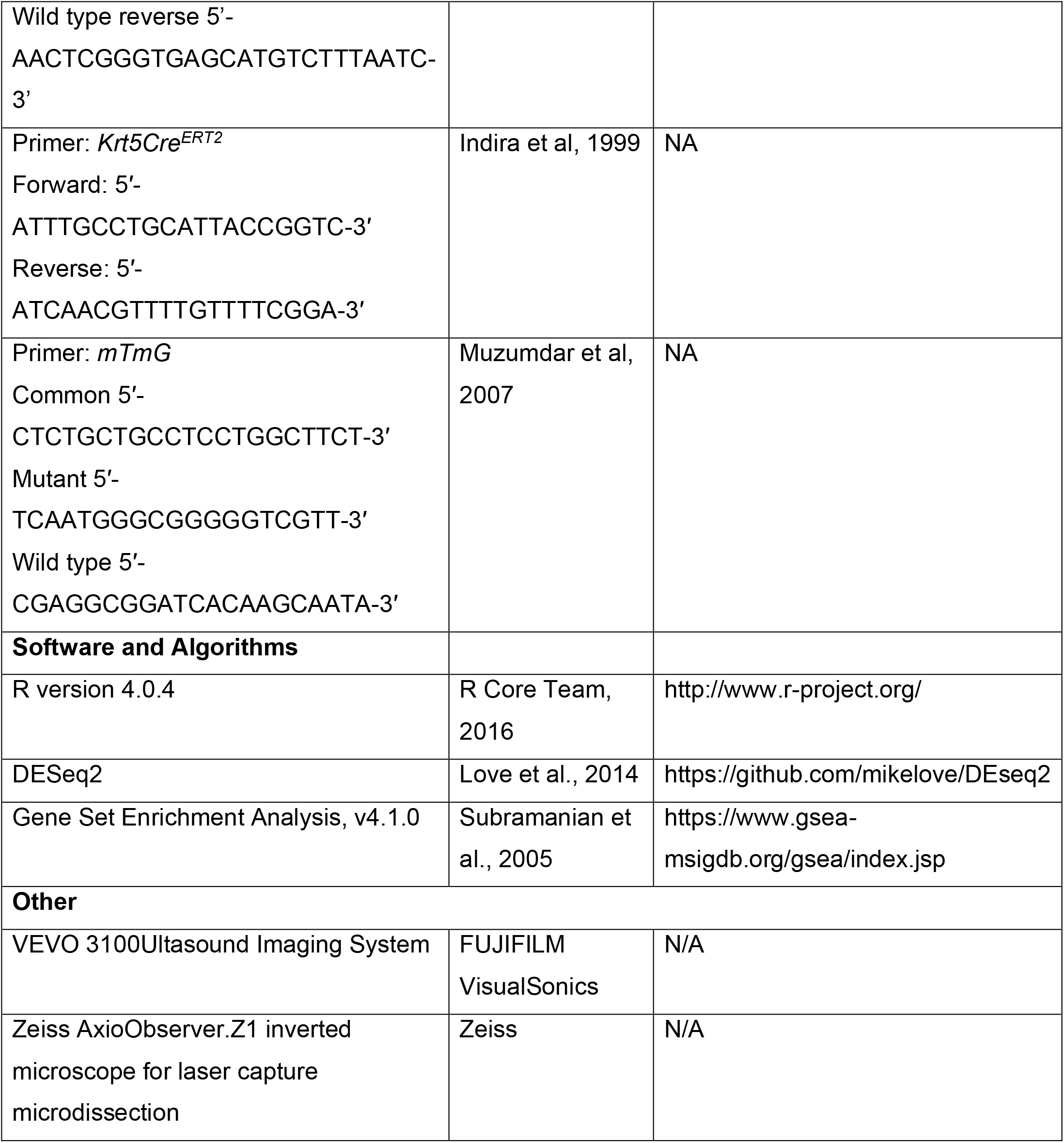

## EXPERIMENTAL MODEL AND SUBJECT DETAILS

### Mice

To generate an inducible *VP16;Pparg* mouse line, a *VP16;Pparg* cDNA was cloned into the AscI site of  the CTV vector (Addgene #15912) via Gibson assembly method to generate pCTV-VP16PPARG gene targeting vector placing a CAG promoter, a STOP cassette (Loxp-Neo-Loxp), the VP16PPARG cDNA and an IRES-EGFP into intron 1 of the ROSA26 gene.  The VP16PPARG is expressed from a synthetic CAG promoter after the removal of the STOP cassette by Cre recombinase. The pCTV-VP16PPARG gene targeting vector was linearized by AloI and electroporated into KV1 (129B6 hybrid) ES cells to generate targeted ES cells with modified ROSA26-CAG-STOP-VP16Pparg allele. The targeted ES cells were injected into the C57BL/6N blastocysts to generate germline chimeras. The male chimeras were bred to wild-type C57BL/6N females to transmit the ROSA26-CAG-STOP-VP16PPARG-IRES-EGFP allele. For conditional tissue-specific expression of the *VP16;Pparg*, the master ROSA26-CAG-STOP-VP16PPARG-IRES-EGFP mouse line was crossed to *Krt5Cre^ERT2^* and induced by Tamoxifen. *mTmGfl/fl* (Gt(ROSA)26Sortm4(ACTB-tdTomato,-EGFP)Luo/J) mice were obtained from Jackson Laboratory (stock #007576). *K5Cre^ERT2^* mice (FVB.Cg-Tg(KRT5-cre/ERT2)2Ipc/JeldJ) were obtained from D. Metzger and P. Chambon. All work with mice was approved by and performed under the regulations of the Columbia University Institutional Animal Care and Use Committee. Animals were housed in the animal facility of Irving Cancer Research Center, Columbia University.

### Human specimens

Bladder tumors were obtained from patients undergoing transurethral resection of tumors (TURBT) at Columbia University Medical Center. All patients gave informed consent under the Institutional Review Board-approved protocols.

## METHOD DETAILS

### Intraperitoneal Tamoxifen induction

Mice (8-12 weeks of age) were injected with tamoxifen (Sigma cat#T5648) dissolved in corn oil, at a dose of 5 mg per 30 g body weight for 5 consecutive days.

### Ultrasound guided Tamoxifen induction

Mice (8-12 weeks of age) were injected with 4-OHT dissolved in DMEMF12/Tween 80 at a dose of 80 ug/mL every other day for two dosages using a VEVO 3100 Ultrasound Imaging System (FUJIFILM VisualSonics, Toronto, Canada) located within the mouse barrier in the Herbert Irving Cancer Center Small Animal Imaging facility.

### BBN Treatment

BBN (0.05%; Sigma cat#B8061-1G) was administered in the water supply daily for 3.5 – 24 weeks to induce bladder cancer. Mice were euthanized at 3.5 or 24 weeks. All bladders were removed and embedded for sectioning and staining.

### Laser Capture Microdissection (LCM)

5 month BBN *K5VP16;Pparg* and control bladders were flash frozen sagitally in OCT and stored at −80°C. The specimen blocks were placed in the Leica CM3050 cryostat chamber for 5-10min to temperature equilibrate. The tissue was sectioned at 5 μm on Arcturus PEN Membrane glass slides (Applied Biosystems cat#LCM0522) and the slides were immediately stored at −80°C. 1 hour prior to LCM, slides were removed from the −80°C and stained with Arcturus HistoGene Staining Solution (ThermoFisher Scientific cat#KIT0415) according to the manufacturer’s protocol. Tissue was then laser captured on a Zeiss AxioObserver.Z1 inverted microscope into AdhesiveCap 200 opaque caps (Zeiss cat#000830-19). 240μL of RLT Lysis Buffer (Qiagen cat#1015750) was then added to the captured tissue. The suspension was then processed for total RNA extraction. Samples with a RIN (regulation identification number) >6.5 were used for RNA-seq. These samples then were sequenced according to the steps listed in the RNA-Sequencing method.

### Single Cell Dissociation

Cold active protease (CAP), was prepared and stored on ice (5mM CaCl2, Sigma cat#21115, 10mg/mL Bacillus Licheniformis protease, Sigma cat#P5459, 12.5U/mL DNAse, Sigma cat#4716728001). Mice were perfused with 20 mL of CAP using a small vein infusion set (Kawasumi cat#D3K2-23G) and two 10 mL syringes per mouse. Bladders were dissected and immediately put in a 60mm x 15mm petri dish (Fisherbrand cat#FB0875713A) containing CAP on ice for 10 min. The bladders were then transferred to HBSS media (ThermoFisher cat#14170-112) containing 1% Bovine Serum Albumin (Sigma cat#A2058) and 0.1% glucose, where they were inverted and the urothelium was manually separated from the stroma. The cell suspension was then collected into a 1.5mL Eppendorf tube on ice and gently triturated until the cells were in a single cell suspension.

### RNA-Sequencing

For homeostasis experiments, *K5VP16;Pparg;mTmG* and *K5;mTmG* single cell suspensions were filtered through a 70μm filter (Fisherbrand cat#22363548) and then sorted on a BD Aria II Cell Sorter using a 130 μm nozzle aperture and 13 psi pressure to collect GFP-positive cells. Cells were then centrifuged at 500 x g for 30 min at 4°C. The supernatant was discarded, and the pellet was processed for total RNA extraction. Samples with a RIN (regulation identification number) >8 were used for RNA-seq. The libraries were prepared using the SMART-Seq^®^ v4 Ultra^®^ Low Input RNA Kit for Sequencing (TaKaRa) followed by Nextera XT (Illumina), both according to manufacturer’s instructions. They were sequenced to a targeted depth of 40M 2×100bp reads on a NovaSeq 6000 (Illumina). Differential expression analysis was performed by reading kallisto counts files into R using the R packages tximport (v.1.10.1) and biomaRt (v.2.34.2),and running DESeq2 (v.1.18) to generate log fold change values and p-values between the two experimental groups. The heatmap and PCA plots were visualized after transforming the counts using VST (variance stabilizing transformation). Gene set analysis by ConsensusPathDB (Kamburov, A. et al. 2013) were used to identify significantly changed pathways.

### Heatmaps

RNA reads were aligned to the mouse reference genome(mm10) using STAR (v2.5.3a). The transcript-levels were then quantified using SALMON (v0.9.1). Count data were extract from SALMON output using Tximport (Bioconductor), and normalized and log2 transformed using DESeq2 (Bioconductor). Heatmaps were generated by ComplexHeatmaps(Gu et al., 2016). Box plots of selected genes were generated by ggplot2(R 3.6.2). Significance was calculated using Mann-Whitney U test. Heatmaps used unsupervised clustering of *VP16;Pparg* mouse tumors and BBN/UPPL mouse tumors across previously established immune gene signatures(Saito et al., 2018). Each gene signature was calculated as the z score of average value of all genes included in the signature. Gene expression of murine tumors and human tumors from the The Cancer Genome Atlas (TCGA; n = 408) was combined using upper quantile normalization and co-clustered using Basal/Luminal marker genes.

### Statistical analysis

All quantitation was performed on at least three independent biological samples, using the ImageJ software. Data presented in box plots are mean values ± s.e.m. Statistical analysis was performed using the R version 4.0.4. In two group comparisons, statistical significance was determined using Mann-Whitney U test, considering a value of p <0.05 as significant. The number of samples used in the experiments is included in figure legends.

### Immunostaining

Bladders were embedded in paraffin and serial sections were generated. For immunohistochemistry, paraffin sections were deparaffinized using HistoClear and rehydrated through a series of Ethanol and 1× PBS washes. Antigen retrieval was performed by boiling slides for 15 min in pH 9 buffer or 30 min in pH 6 buffer. Primary antibodies in 1% horse serum were incubated overnight at 4 °C. The next day, slides were washed with PBST three times for 10 min each and secondary antibodies were applied for 90 minutes at room temperature. DAPI (4′,6-diamidino-2-phenylindole) was either applied as part of the secondary antibodies cocktail or for 10 min, for nuclear staining, and then the slides were sealed with coverslips.

### Fluorescent microscopy

Zeiss Axiovert 200M microscope with Zeiss Apotome were used to collect immunofluorescent images. Bright-field images were collected using a Nikon Eclipse TE200 microscope. Data was analyzed using the Fiji package of ImageJ.

## Lead Contact

Further information and requests for resources and reagents should be directed to and will be fulfilled by the Lead Contact, Cathy Mendelsohn.

## Data and Code Availability

Original and source data for RNAseq in the paper are available at GEO: GSE172656

## Acknowledgments

We thank Ronald M. Evans for the *VP16;Pparg* vector; Daniel Metzger for the *Krt5^CreERT2^* line; Renu Virk for pathological evaluation of the tumors; Luis A. Pina Martina for obtaining human tumors; Lauren Shuman and David Degraff for assistance with laser capture microdissection; Mathieu Rouanne for critical reading of the manuscript; Chang Liu for discussions of experimental design; Yinglu Li for assistance in sequencing analysis; Jonathan Ruhl for genotyping. This work was supported by: NIDDK R01 DK095044 (C.L.M), NIDDK U01 DK094530 (C.L.M), TJMCU508-5926-URO, and T32 Training Grant DK07328 (T.T./T.X.). This research used the resources of the Herbert Irving Comprehensive Cancer Center Genetically Modified Mouse Model Shared Resources, Flow Cytometry Shared Resources, Confocal and Specialized Microscopy Shared Resources, Molecular Pathology Shared Resources, Genomics and High Throughput Screening Shared Resources, and Oncology Precision Therapeutics and Imaging Core funded in part through Center Grant P30CA013696.

## Author Contributions

T.T. designed experiments and analyzed the phenotypes in *K5VP16;Pparg* mutant mice, prepared figures, and wrote the paper. T.X. assisted in designing experiments and analyzed the phenotypes in *K5VP16;Pparg* mutant mice, prepared figures, and wrote the paper. M.Z. performed the TCGA and immune signatures analyses. W.Y.K and C.L. helped interpret results. X.C. assisted in interpreting sequencing results. H.K. assisted in laser capture microdissection. E.B. performed immunostaining. C.S.L. created the *VP16;Pparg* mouse construct. S.E.W. performed pathological grading of tumors. J.M.M. provided human tumor samples. C.L.M helped design experiments, interpret results, and wrote the paper.

## Notes

### Competing Interest Statement

The authors have declared no competing interest.

