## Supplementary data for "*Pparg* drives luminal differentiation and luminal tumor formation in the urothelium"

Supplementary data related to Figure 1.

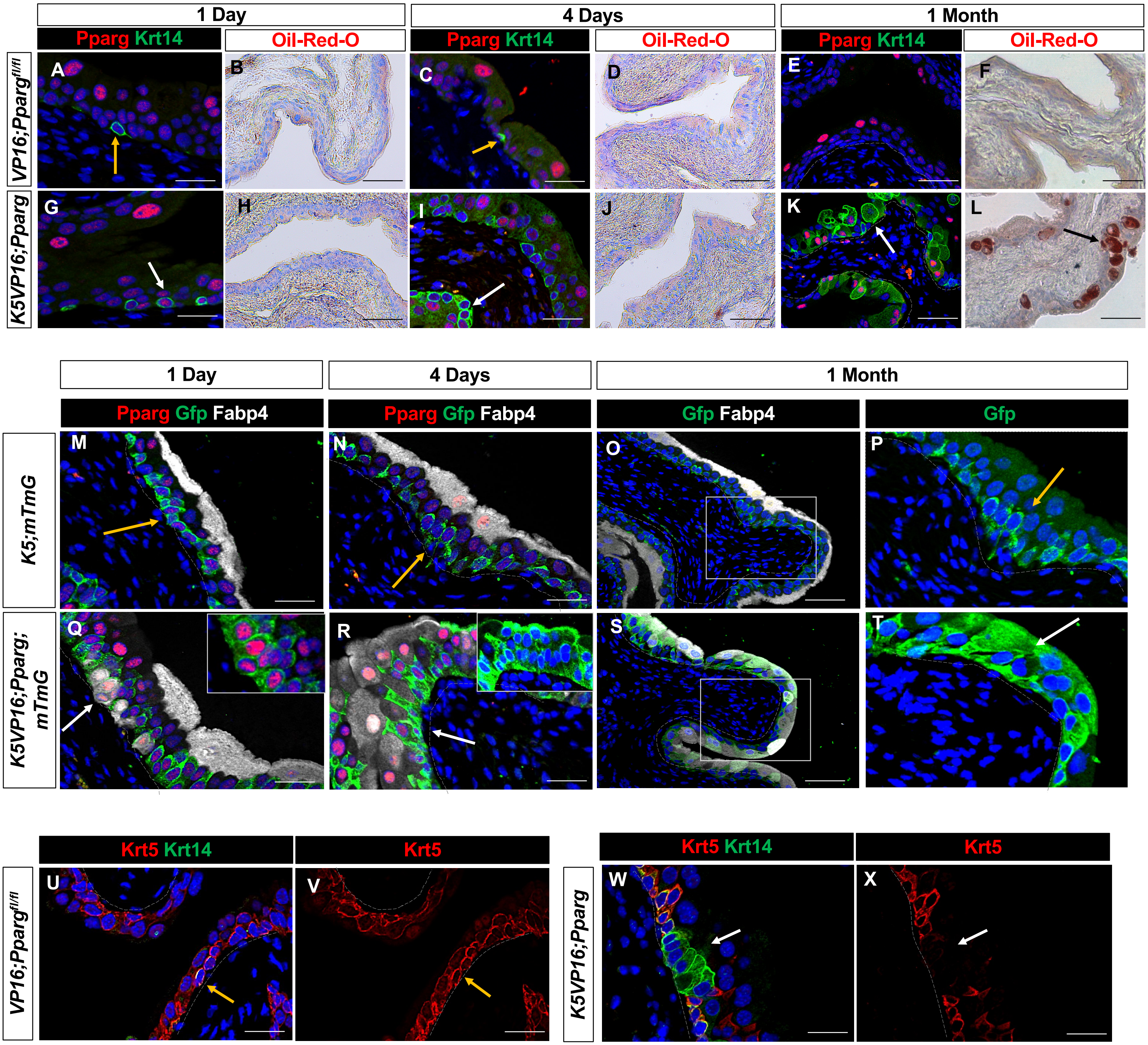


Y

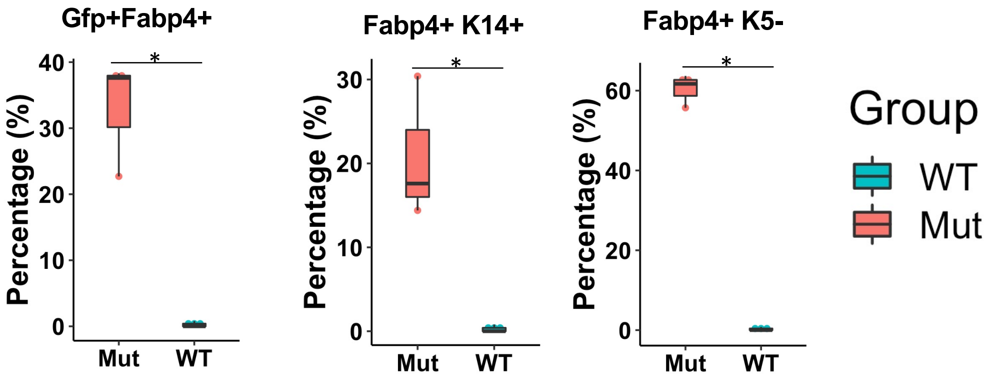


Z

| RA Targets | Tam + 4 Days | |
|---|---|---|
| Gene | FC | Pvalue |
| Aldh1a3 | 1.851856 | 0.005908 |
| Crabp2 | 13.009396 | 0.000322 |
| Rbp4 | 2.230968 | 0.000612 |
| Rdh11 | 1.532477 | 0.000320 |
| Stra6 | 3.174773 | 0.036267 |
| Dhrs9 | 2.526637 | 0.017307 |
| Dgat2 | 1.678491 | 0.000150 |
| Ttr | 8.088511 | 4.01E-11 |

Supplementary data related to Fig. 2

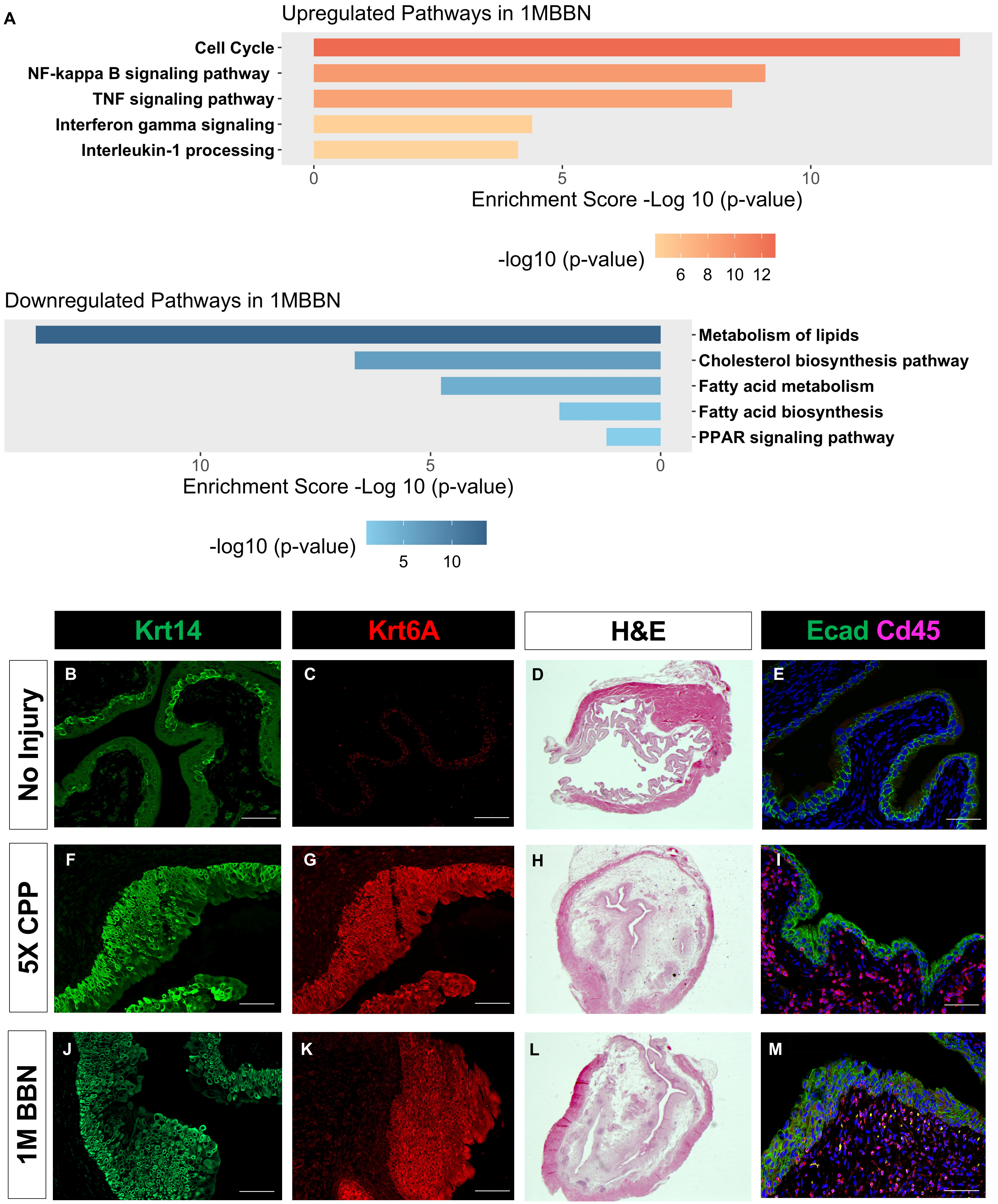

Supplementary data related to Figure 3

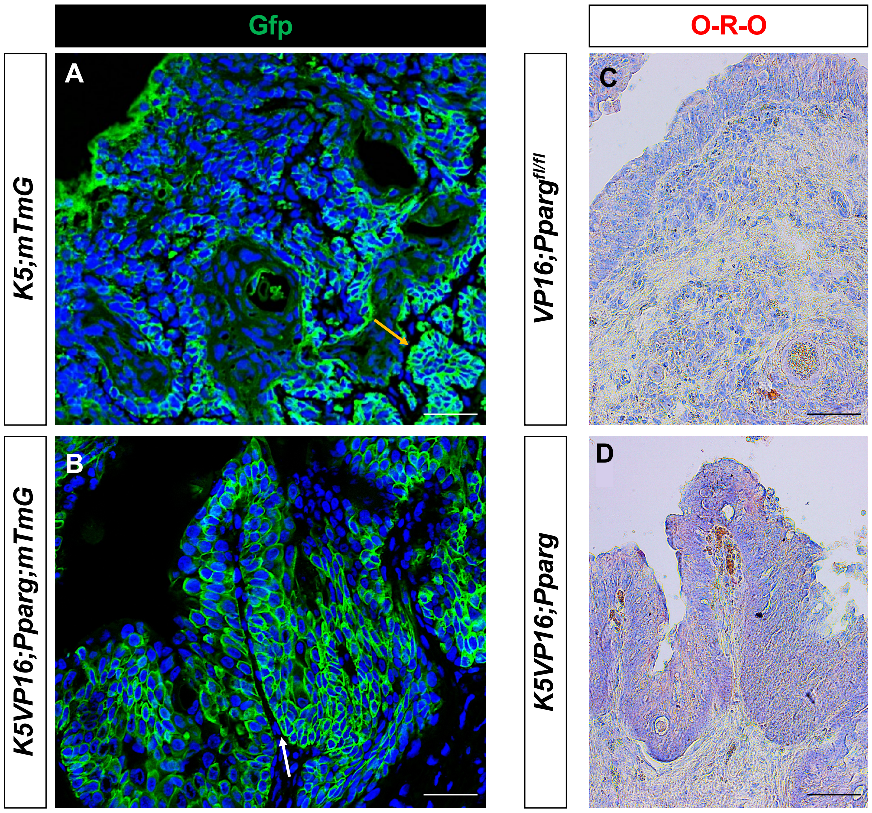


E

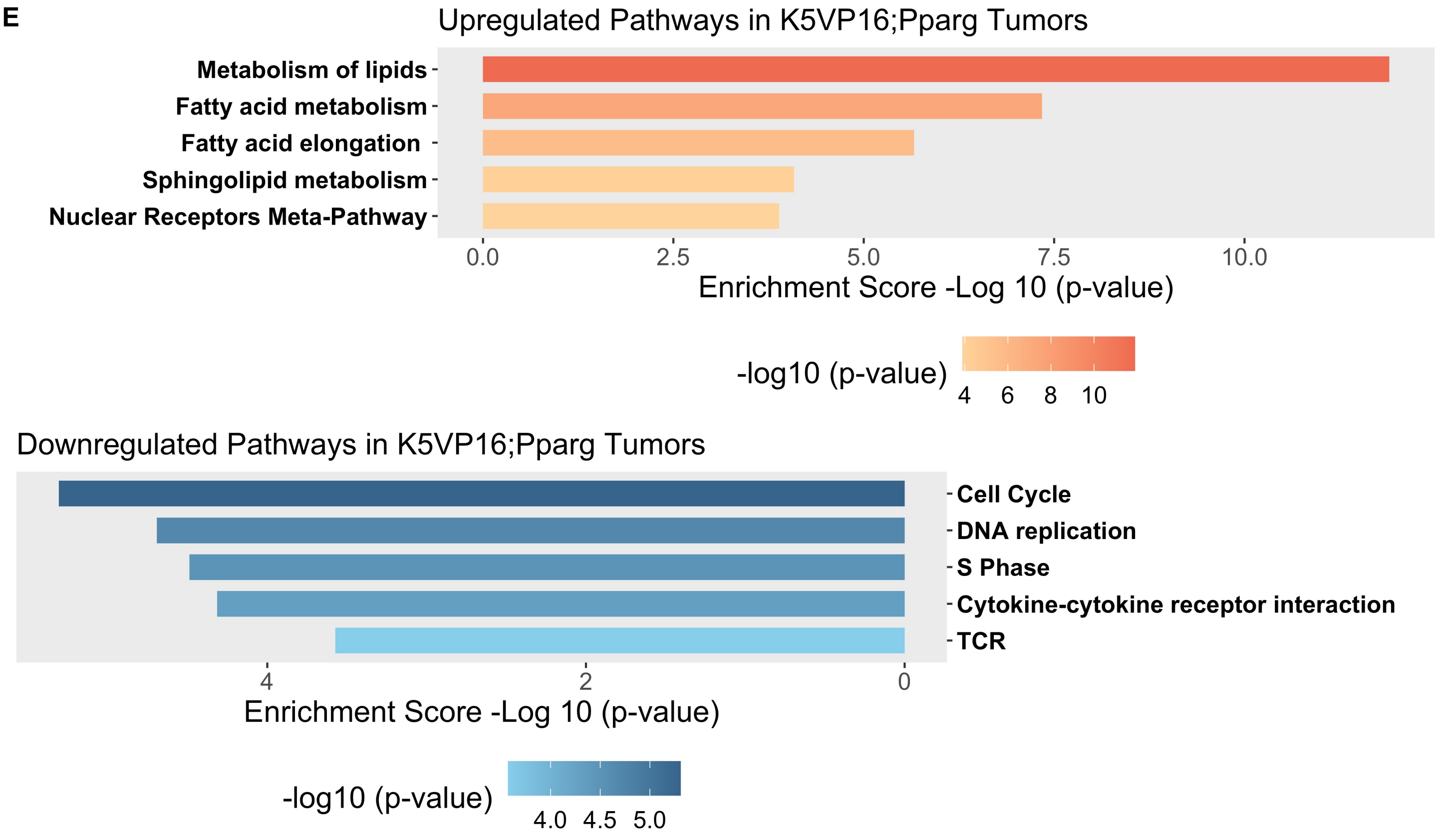

Supplementary data related to Fig. 6

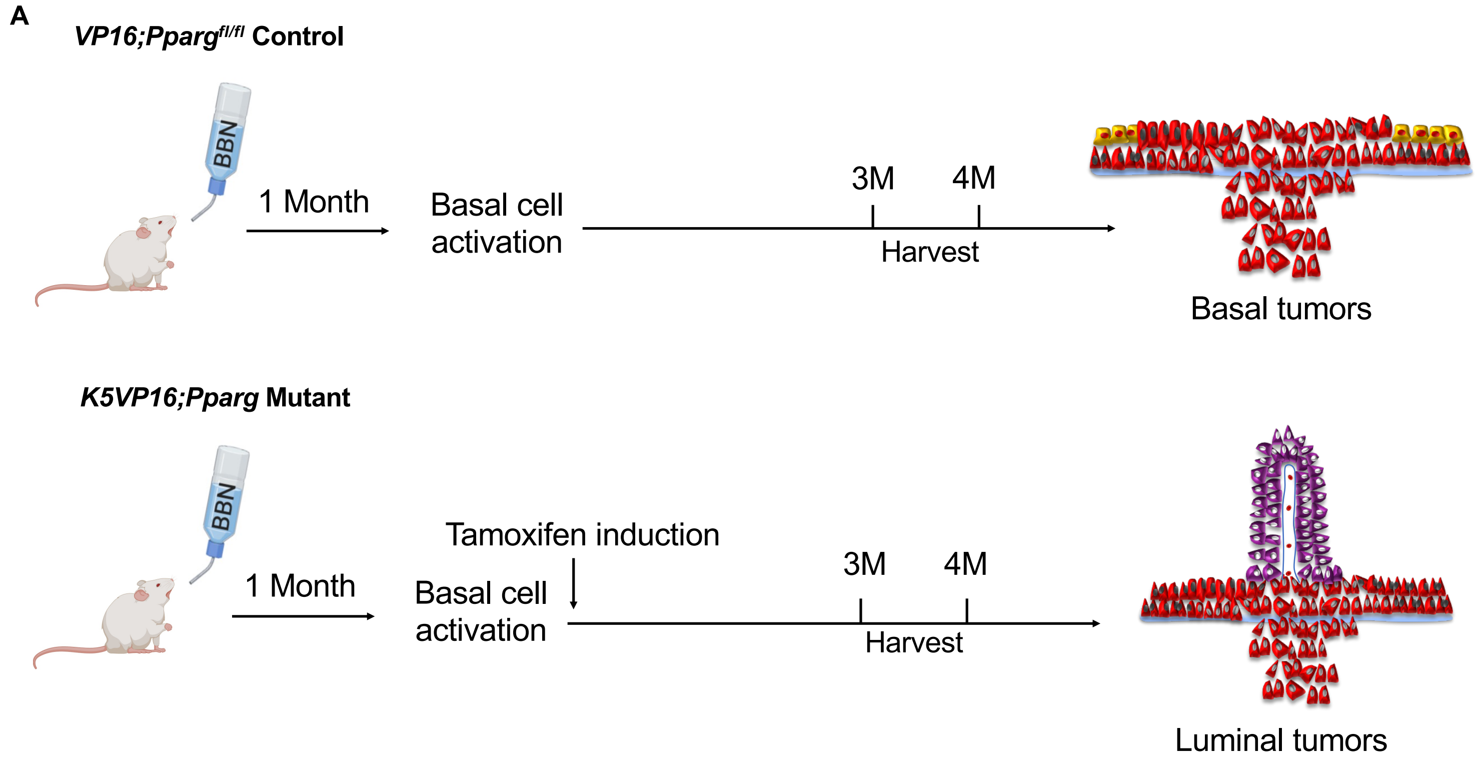
